# What abundance correlations actually measure in stochastic ecological communities

**DOI:** 10.64898/2026.08.12.744396

**Authors:** Akiva Goldberg, Nadav M. Shnerb

## Abstract

Abundance correlations cannot reveal ecological interactions without an assumption about the covariance of environmental noise. A natural biological expectation is that similar species respond similarly to environmental fluctuations, generating positive correlations. Yet the same species also tend to overlap more strongly in resource use and therefore compete more intensely, generating negative correlations. The simplest plausible benchmark is thus to take environmental-response correlations proportional to niche overlap. We show that, under this assumption and across a broad class of stochastic community models, the two effects cancel exactly: equal-time abundance correlations vanish, independently of interaction strength, heterogeneity, and system size. Away from this matched point, the observed correlations measure primarily the mismatch between shared environmental response and competition, rather than the interaction matrix itself. Correlations can recover information about niche overlap when competitive feedback is delayed relative to environmental forcing, but the inference then depends on a resource-response timescale that is generally not determined by the abundance time series alone. When stochasticity enters through the mechanism that generates similarity itself—for example, through fluctuating shared resources—nonzero correlations may persist, but they reflect yield–depletion mismatch rather than niche overlap. Abundance correlations therefore report how environmental variability reaches the community at least as much as they report who competes with whom.

## I. INTRODUCTION

Time-resolved community data have become abundant, and with them the hope of reading the structure of a community off the fluctuations of its species. Where two species rise and fall together, one is inclined to suspect that they are alike, where they rise and fall in opposition, that they compete. Abundance correlations have accordingly been pressed into service as proxies for ecological interaction, and as time-resolved community data become more widely available, in microbial systems above all, these inferences are attracting growing interest [1–5].

Such an inference is not straightforward. Two closely related species are similar in two respects at once, and the two work against each other. They are similar in physiology, and therefore respond alike to the environment: a warm year favours both, a dry one harms both, and their abundances are drawn together. On the other hand, they are also similar in their resource use, and therefore compete: when one prospers it consumes what the other needs, so their abundances are driven apart. Relatedness thus generates a positive correlation and a negative one simultaneously, and the sign of what is finally observed is not given in advance. Which effect prevails?

The question is quantitative, and it is exactly the kind of question a model of community dynamics ought to be able to answer. In the standard treatments, environmental variability is introduced by adding stochastic terms to Lotka–Volterra equations [6–8], and the correlation structure of that noise is then imposed from outside. But if the interaction matrix is to be read as niche overlap [9, 10], the two cannot be independent: the biological similarity that sets the competition must also be reflected, at least in part, in the correlation of the environmental responses.

In this paper we therefore try to constrain the noise structure using a biologically motivated benchmark and ask what follows. We work with two models, using them as instruments rather than as subjects. The Lotka–Volterra description is generic and transparent [11], and in it the balance can be settled exactly. But it has no resources in it, and therefore no account of two things that turn out to matter: that competition, being mediated by a shared resource pool, does not act instantaneously, and that what a consumer takes from a resource and what it gains from it need not be proportional. For these we turn to the consumer–resource description [12], in which both are explicit.

The central difficulty is that there is no obvious way to determine how strongly correlations in environmental forcing should track competitive similarity. A natural benchmark nevertheless suggests itself: the correlations between species’ environmental responses may be taken to be proportional to the biological similarity encoded by their interactions. This prescription is quite general and includes, as a special case, the standard consumer–resource setting in which yield and depletion are proportional. It leads to a surprising conclusion: at the matched point, where environmental-response similarity coincides with interaction similarity, the effects of shared forcing and competition cancel exactly, and the abundance correlations vanish.

Under this benchmark, the cancellation is not merely a leading-order effect: the full nonlinear stationary distribution factorizes into independent single-species Gamma laws at arbitrary noise strength. Consequently, all equal-time interspecific correlations vanish, even though the interactions remain present in the dynamics. The resulting Gamma abundance distributions reproduce some of the macroecological patterns emphasized by Grilli [13], Azaele *et al*. [14], but persist here for a broad class of interaction matrices. The corresponding factorized stationary state was established previously for reciprocal interactions by Camacho-Mateu *et al*. [15]. Appendix C extends this result to nonreciprocal dynamics, where the same stationary distribution coexists with a nonvanishing probability current.

This observation calls into question the use of equal-time correlations to infer competition. What abundance correlations measure, in consequence, is not competition but the residue, the extent to which the environmental forcing fails to match the interactions. That residue, we show, carries no information about the interactions.

There are two exceptions where the correlations provide information about interactions, and both are instructive. The first is a matter of timing. Competition is not a force that one species exerts on another directly, it is transmitted through the resources they share, and its arrival is delayed by the time the resource pool takes to respond. The shared environmental response suffers no such delay. When the resources are slow, therefore, the cancellation fails and the correlations do track niche overlap, decaying with phylogenetic distance in the manner that is observed by Sireci *et al*. [2]. Within the models we analyze, this is the one circumstance in which the conventional reading of a correlation matrix is sound. It is not, however, a circumstance the data announce: it depends on a resource timescale that consumer-abundance measurements do not reveal.

The second exception arises when the environment acts on the resources rather than on the consumers [4, 16], and requires no assumption about how species respond to it: the resources fluctuate, and the consumers inherit whatever correlation their shared reliance on those resources implies. Here, nonzero correlations may appear only if yield and depletion are misaligned, if what a species removes from a resource is not proportional to what it gains. The signal is then real and strong, but it is a signal about the misalignment, not about the competition.

In real communities, the route through which environmental variability enters the dynamics is generally unknown. As the two constructions above illustrate, the same interaction matrix can therefore be associated with very different abundance-correlation patterns. Equal-time correlations alone cannot distinguish among these possibilities and thus should not, in general, be interpreted as a direct map of species interactions.

The paper is organized as follows. Section II sets out the two descriptions, the Lotka– Volterra and the consumer–resource, and the mapping between them. Section III shows that equal-time correlations cannot identify the interaction matrix without an assumption about the environmental noise. Section IV adopts the simplest such assumption, that the noise correlation matches the niche overlap, and shows that the correlations then vanish identically, so that what they measure away from this point is the response–competition mismatch rather than the interactions. Sections V and VI return the resources to the model and separate the two channels through which the environment acts, giving the two circumstances in which a signal does survive, retarded competition and resource-mediated noise. We conclude by discussing what this implies for the modelling of environmental stochasticity, and for inference from community time series.

## II. TWO LEVELS OF DESCRIPTION: WHAT THE REDUCTION PRESERVES, AND WHAT IT DESTROYS

Throughout this paper we implement two standard models of community dynamics: the consumer-resource (CR) model and the Lotka-Volterra (LV) model. These are not competing alternatives but two levels of description of the same interacting system. The CR description retains the mechanistic distinction between the depletion a consumer imposes on a resource and the benefit it derives from it, while the LV description emerges from it, in appropriate limits, as an effective coarse-grained dynamics. Although we expect our conclusions to hold more broadly, this pair provides an especially transparent framework, and the relation between the two levels will carry the argument of the paper.

Consider a community of *S* consumer species competing for *Q* resources. The abundance of consumer *i* is *ni*(*t*) and the biomass of resource *k* is *Rk*(*t*). Consumer growth is driven by resource uptake, while each resource grows logistically in the absence of consumers and is depleted through consumption:

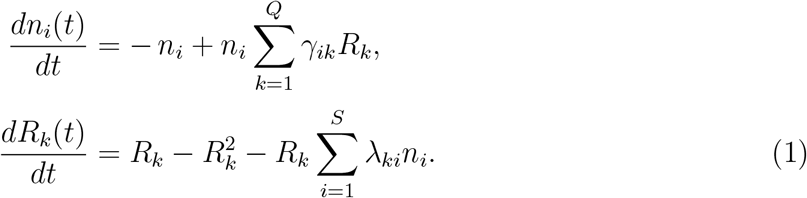

Here *γ*_*ik*_, the (*i, k*) entry of the *S* × *Q* matrix Γ, quantifies *yield* : the increase in the growth rate of consumer *i* per unit of resource *k. λ*_*ki*_, the (*k, i*) entry of the *Q* × *S* matrix Λ, quantifies *depletion*: the per-capita rate at which consumer *i* removes resource *k*.

### A. Yield and depletion are distinct

The essential feature of the formulation (1) is that it keeps Γ and Λ apart. In traditional consumer-resource models [12] they are usually tied together: yield and depletion are taken proportional at the level of the individual consumer-resource interaction,

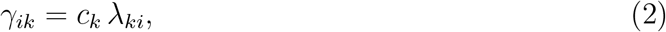

with a conversion factor *ck* that depends only on the resource. Consumption and growth are thereby identified up to a fixed (per resource) efficiency.

We relax this assumption and allow benefit and depletion to be misaligned. Such a decoupling can arise from differences in conversion efficiency, from wasteful or destructive uptake, from metabolic by-products and cross-feeding, and from related processes [17–24]. Furthermore, recent work suggests that the yield-to-depletion ratio is not fixed, but varies across conditions, resources and taxa [25–27].

### B. The reduction to Lotka-Volterra

To obtain an effective Lotka-Volterra description one integrates out the dynamics of the resources. Setting *dRk/dt* = 0 gives the quasi-steady resource levels

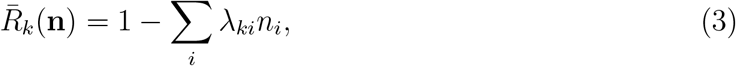

which, substituted back into the consumer equation, yield

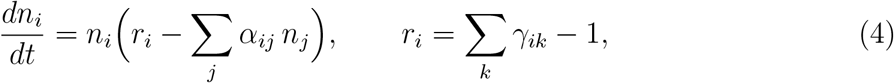

with the effective interaction matrix

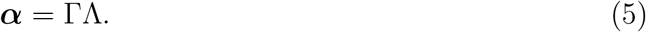

The assumption *dRk/dt* = 0 is justified in two closely related limits: when the resource dynamics is much faster than the consumer dynamics, so that the resources may be treated as fast variables, and when the system is considered close to equilibrium. In Sec. V we shall reinstate the resource timescale *τ*_*R*_ explicitly and control this limit rather than assume it.

In what follows we adopt the normalization *α*_*ii*_ = 1 and *ri* = 1, so that Eq. (4) takes the familiar form

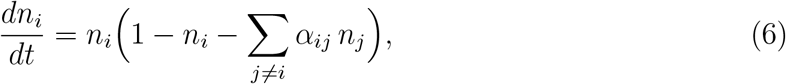

which is the form in which competitive and symmetric Lotka–Volterra communities are customarily analyzed [11, 16, 28–30]. In the numerical work the condition is imposed when the traits are drawn, by normalizing the rows of Γ to a fixed sum, which makes *ri* = _*k*_ *γ*_*ik*_ − 1 = 1 the same for every species. To pick Γ and Λ pairs that satisfy both this constraint and reproduce a prescribed ***α*** = ΓΛ, we implemented a constrained bilinear least-squares technique, see Appendix A.

As explained in Appendix C, our results are completely general and hold for arbitrary choices of the self-interaction coefficients *α*_*ii*_ and intrinsic growth rates *ri*. The present formulation, with identical *α*_*ii*_ and *ri*, is adopted solely for convenience of presentation.

### C. Niche overlap

The matrix ***α*** carries the entire biological content of the Lotka-Volterra description. With the diagonal normalized to unity, *α*_*ij*_ is naturally read as the effective *niche overlap* of consumers *i* and *j* [9, 10, 31]. When it vanishes the two consumers exploit distinct resources and do not constrain one another’s growth, a value close to one corresponds to nearly maximal overlap.

Once yield and depletion are allowed to differ, the product ***α*** = ΓΛ is generically *not symmetric*. The overlap of a pair is then carried by the symmetric part ***α***_*S*_ = (***α*** + ***α***^*T*^)*/*2, while the antisymmetric part ***α***_*A*_ carries a second, distinct quantity, the directional asymmetry of the interaction, the extent to which *i* affects *j* more than *j* affects *i*. The antisymmetric part of ***α*** has to do with the fitness differences between the species [10].

### D. When the reduction is legitimate, and what it discards

The imposition of *dRk/dt* = 0 is legitimate for the *static* properties of the community: the location of the fixed point, and any question posed in terms of the equilibrium alone, need only the effective matrix ***α*** = ΓΛ. It ceases to be legitimate once the dynamics matters. Away from equilibrium the yield Γ and the depletion Λ, which the reduction melts into a single product, reassert themselves separately, and communities sharing an identical nicheoverlap matrix can behave in opposite ways [16].

The mechanism is visible directly in Eq. (4): if the resources are displaced from quasi-steady by *δRk*, the growth rate of consumer *i* is displaced immediately by

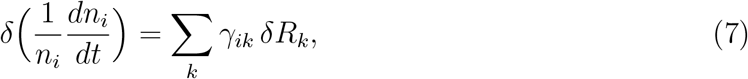

which involves the yield profile of *i* but not Λ. As explained below this separation has to do with the behavior of abundance correlations.

## III. ABUNDANCE CORRELATIONS DO NOT IDENTIFY THE INTERACTIONS WITHOUT AN ASSUMPTION ON THE NOISE

Suppose environmental variability is introduced at the Lotka–Volterra level in the standard way [6–8], as a random term in the growth rate:

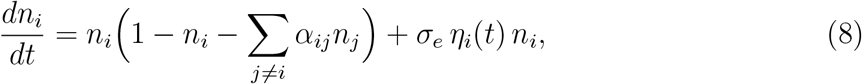

with 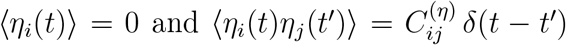. Both matrices are normalized to a unit diagonal: *α*_*ii*_ = 1 and 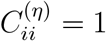, the amplitude of the stochastic forcing being *σ*_*e*_. Simulations below use the Stratonovich convention (see [16]), but the qualitative picture does not depend on this choice as explained in Appendix C.

Let us assume that the deterministic part of Eq. (4) supports a stable coexistence state. 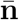 is the vector of population densities at equilibrium, which satisfies 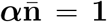, and 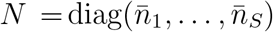 is the corresponding diagonal matrix. Linearizing Eq. (4) about this fixed point, the deviations 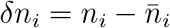 obey a multivariate Ornstein–Uhlenbeck equation,

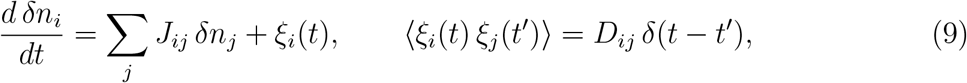

with drift matrix and noise covariance

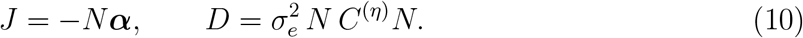

The stationary covariance ∑_*ij*_ = ⟨*δni δnj*⟩ therefore satisfies the Lyapunov equation

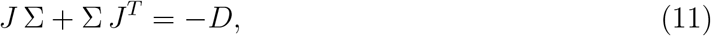

and the normalized abundance correlations are related to ∑ via

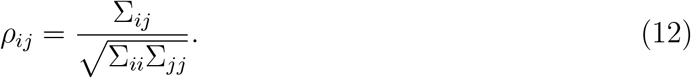

In what follows, *abundance correlation* means the equal-time correlation between the temporal fluctuations of two species about a stationary state, measured within a single community.

Equation (11) relates three objects: the drift *J*, which carries the interactions, the covariance ∑, which is what the abundance correlation data provide, and the diffusion *D*, which carries the stochastic environmental forcing. One measurement, of ∑, cannot determine two unknowns. Any attempt to read ***α*** off ∑ must therefore say, in advance, what *C*^(*η*)^ is.

This redundancy is illustrated in Fig. 1. Here we took a community of six species and a 6 ×6 correlation matrix *C*^(*η*)^ for the stochasticity, and obtained the covariance matrix ∑. We then constructed two further communities, one bearing no relation to the “true” community and one in which the similarity relations between the species are exactly reversed, and by suitably tailoring *C*^(*η*)^ for each of them we reproduced exactly the same abundance correlation matrix. The lesson is clear: the interactions cannot be inferred from the correlations without saying something about the covariance of the external stochasticity.

**FIG. 1.**
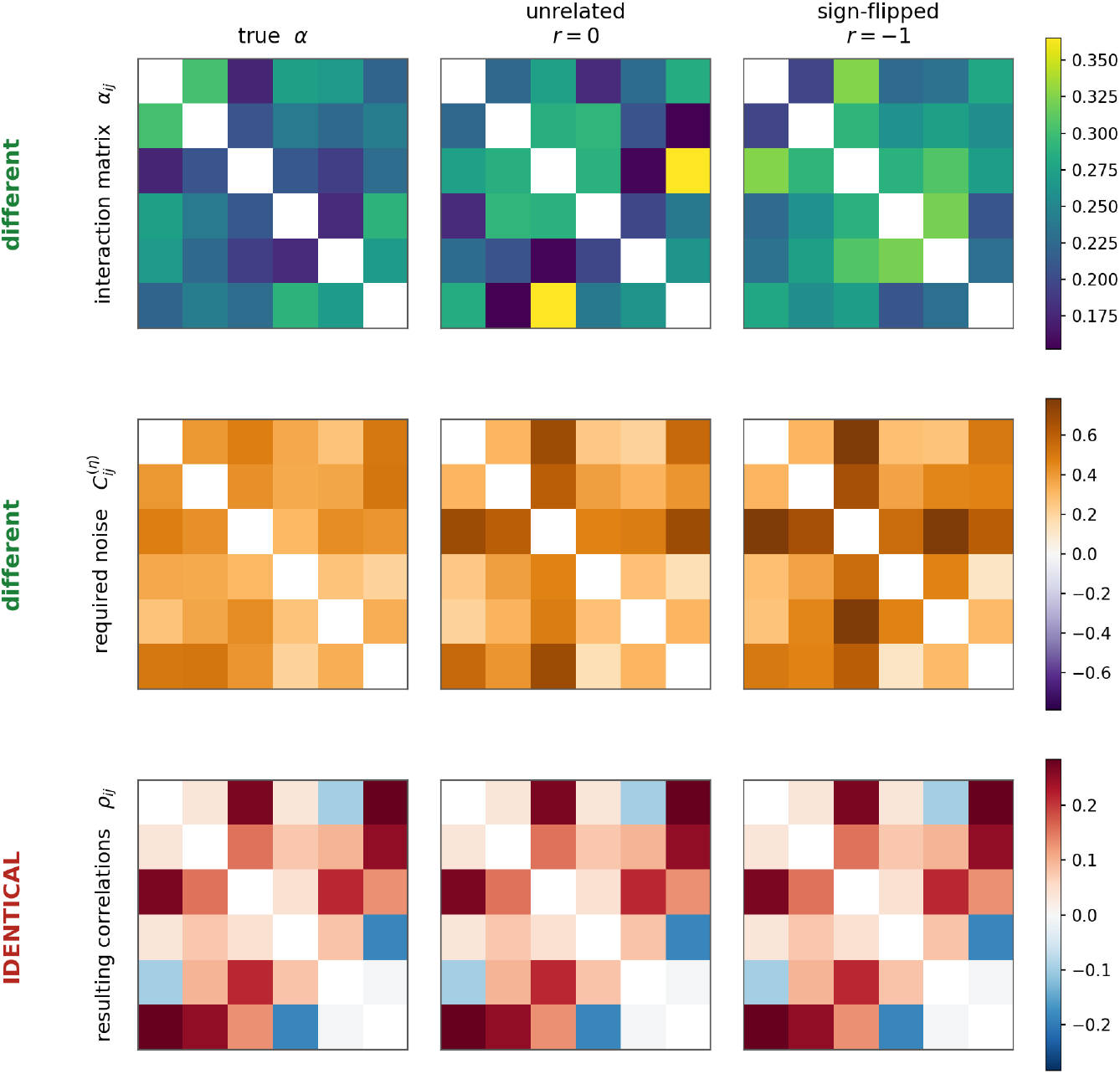
The interaction matrix cannot be recovered from equal-time correlations. The dynamics of three different communities of *S* = 6 species, each with its own interaction matrix (top row), were simulated. All three matrices are legitimate, with positive off-diagonal entries in the range [0.15, 0.37] and a stable coexistence fixed point, and all three yield exactly the same abundance correlations (bottom row). To achieve this, the noise covariance matrix *C*^(*η*)^ (middle row) was obtained from Eq. (4), given ***α*** and ∑^*^. The first two interaction matrices (left and middle columns) were picked at random, with the same mean overlap but no relation pair by pair. In the third community (right column) the heterogeneity is *reversed* with respect to the first: pairs that in reality overlap most strongly are assigned the weakest overlap, and conversely, yet by tailoring the correct *C*^(*η*)^ one obtains the same correlation matrix. The marginal variances, which are not part of the correlation data, being chosen to render each reconstructed *C*^(*η*)^ of unit diagonal.

The algorithm by which we constructed the required *C*^(*η*)^ matrices proceeds as follows. Suppose ∑^*^ has been measured with perfect precision, and take any stable candidate interaction matrix. It fixes its own equilibrium, 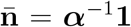, and hence both *N* and *J* = −*N* ***α***. Eq. (4) then returns, in one step, the exact noise that this candidate requires:

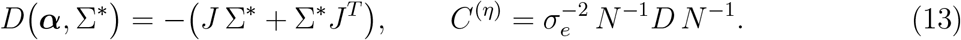

The impossibility of recovering the deterministic part of the dynamics from the correlations of the fluctuations about a fixed point is, in general, well known [32, 33]. In our particular case, however, one must still verify that the *C*^(*η*)^ demanded by Eq. (4) is a genuine correlation matrix: symmetric, positive semidefinite, and with unit entries on the diagonal. Neither of the last two is guaranteed by the construction. We have checked it numerically, and as Fig. 1 shows there is no difficulty: the same abundance correlations were reproduced exactly by three entirely different interaction matrices. Further details, and the check that the reconstructed noise covariance is positive semi-definite with unit diagonal, are given in Appendix D.

In summary, no information about the interactions can be obtained without knowing, or assuming, something about the covariance of the stochasticity. It is hard to conceive of a way to obtain this information from direct measurement, and we must therefore resort to some reasonable biological assumptions.

## IV. THE BIOLOGICALLY GROUNDED NOISE ANNIHILATES THE SIGNAL: CORRELATIONS MEASURE THE MISMATCH

What, then, can be said about *C*^(*η*)^ on biological grounds?

### A. A biologically motivated matched benchmark

The natural answer is that closely related species should experience more strongly correlated environmental forcing. Two independent arguments support it.

A. *Shared physiology*. Closely related species are built alike. They share metabolic pathways, thermal tolerances, water and nutrient requirements, and so respond in similar ways to variation in temperature, humidity, pH, or any other external driver.
B. *Shared resources*. Closely related species also tend to consume the same resources. When the supply of a resource fluctuates, the species that depend on it are affected together, and again the covariance of the forcing they experience should reflect how much of their resource use they hold in common, that is, their niche overlap.

At the Lotka–Volterra level these two arguments are indistinguishable: both say only that species resembling one another are forced alike, and since the sole object available to express this is ***α***, both arguments point to *C*^(*η*)^ tracking ***α***.

Since *C*^(*η*)^ is symmetric it can track only 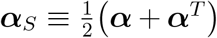, which is the part interpreted as niche overlap between the species [9, 10, 31]. The simplest parameter-free choice, and the matched benchmark we adopt, is therefore,

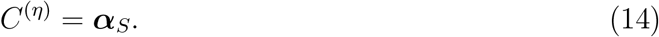

We assume throughout that the interaction matrix ***α*** admits a stable coexistence equilibrium with strictly positive abundances. When ***α*** is symmetric, this is equivalent to ***α*** being positive definite, and therefore ***α*** itself is a valid correlation matrix. When ***α*** is asymmetric, its symmetric part, may nevertheless remain positive definite. In this case we continue to adopt the matched-noise construction of Eq. 14. These are the cases considered throughout the present work.

A further assumption is that the diagonal entries of *C*^(*η*)^ and of *α* are equal to one. We make this assumption solely for convenience of presentation. The corresponding generalization is given in Appendix C 3.

### B. The mismatch is the source of abundance correlations

To determine what abundance correlations measure, both at and away from the matched benchmark, let us define the *response–competition mismatch*

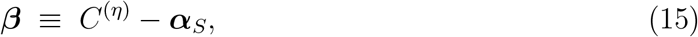

the extent to which the correlation structure of the environmental forcing fails to track the niche overlap. As we now show, this mismatch is the source of every departure of the abundance covariance from its diagonal baseline.

To do that we substitute the ansatz

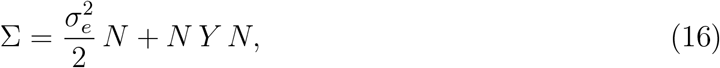

where *Y* is an arbitrary matrix, into Eq. (4). All terms containing ***α*** alone cancel identically, and what remains is

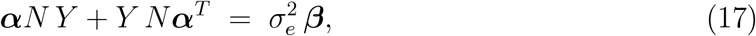

or, more compactly,

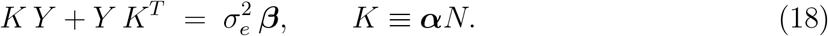

Equation (17) is the organizing result of this paper. Exact within the linearized weak-noise description, and assuming no symmetry of ***α***, it states that every departure of ∑ from the diagonal baseline 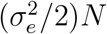 is generated by the single term 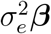.

The mechanism behind this result is transparent already for two species, where Eq. (4) can be solved in closed form:

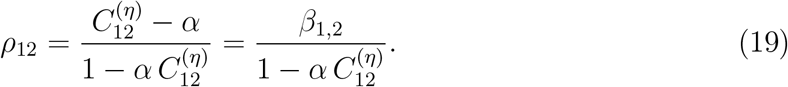

Competition and shared environmental response pull the two abundances in opposite directions, the first anticorrelating them, the second synchronizing them, and the two effects cancel *exactly* when the noise correlation equals the niche overlap, as demonstrated in Figure 2.

**FIG. 2.**
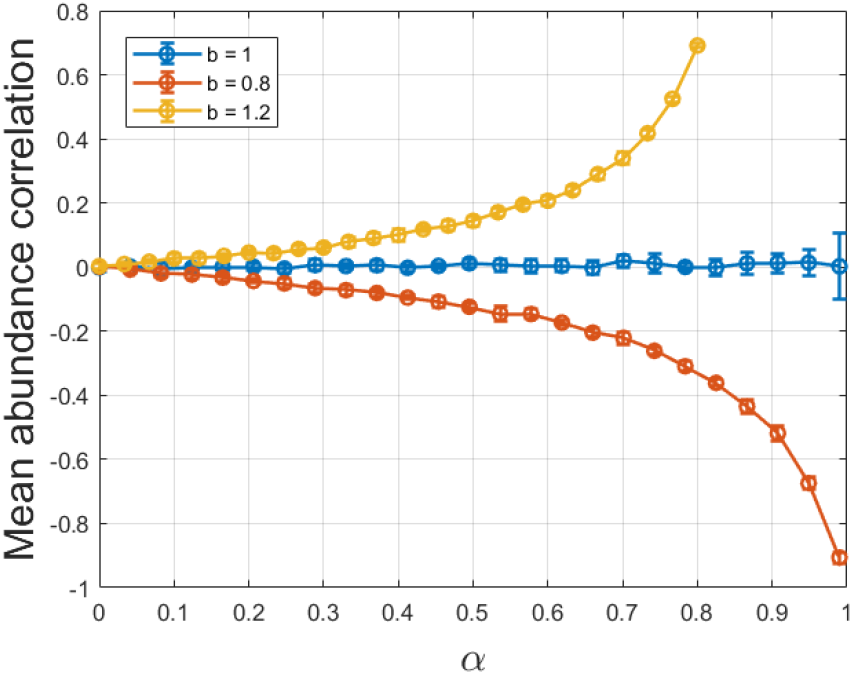
Two species: the abundance correlation is set by the mismatch, not by the niche overlap. Stochastic Lotka–Volterra dynamics, Eq. (4), for *S* = 2 with symmetric competition *α*_12_ = *α*_21_ = *α* and a noise correlation 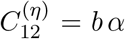. Solid curves are the exact result, Eq. (4), which for this choice of noise reduces to *ρ*_12_ = (*b* − 1)*α/*(1 − *bα*^2^): the correlation is controlled entirely by the mismatch *β*_12_ = (*b* − 1)*α*, and the niche overlap enters only through the propagator in the denominator. Symbols are direct Stratonovich simulations of Eq. (4) with *σ*_*e*_ = 0.05. At the biologically grounded point *b* = 1 the correlation vanishes identically for *every* value of the niche overlap: competition anticorrelates the two abundances and the shared environmental response synchronizes them, and the two effects cancel term by term. Away from that point the correlation is positive when the shared response dominates (*b >* 1) and negative when competition dominates (*b <* 1).

### C. Exact cancellation at the matched point

Equation (17) is linear in *Y*, and the stability of the fixed point guarantees that its solution is unique (see Appendix B 2). If ***β*** = 0 then *Y* = 0 solves the equation, and by uniqueness it is the only solution. The abundance covariance is exactly diagonal and all interspecific correlations vanish identically for any number of species, any degree of heterogeneity, and any degree of asymmetry. Therefore,

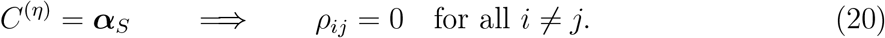

Within the linearized weak-noise description, this cancellation is exact and not merely leading order in ***β***. The derivation and the uniqueness of the solution are given in Appendix B.

At the matched point itself the statement is in fact stronger. There the cancellation is exact and non-perturbative in the noise strength: the full nonlinear stationary distribution factorizes into independent single-species Gamma laws, so that not only the covariance but every equal-time interspecific dependence vanishes, and it does so even when the interactions are non-reciprocal (Appendix C). We nonetheless retain the linearized Lyapunov description throughout, because the object of interest is not the matched point in isolation but its neighbourhood, how the correlations turn on with the mismatch ***β***, for which the Lyapunov equation is the transparent tool.

### D. Near the matched point, correlations reflect the mismatch rather than pairwise interaction heterogeneity

Let us consider a community with weak interaction heterogeneity, i.e.,

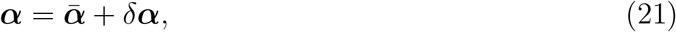

where 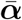 is homogeneous, unit diagonal, all off-diagonal entries equal to the mean overlap *µ*, the average of the off-diagonal entries of ***α***, while *δ****α*** carries the pairwise heterogeneity, the quantity one wishes to infer.

Because the source term in Eq. (4) is *O*(***β***), so is *Y*, hence *δ****α***, appearing only multiplied by *Y*, contributes at *O*(*δ****αβ***). To the leading order (see Appendix B 3) jointly in the mismatch and in the heterogeneity about the homogeneous community, therefore,

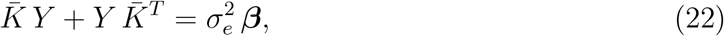

where 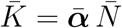 is built from the *homogeneous* interaction matrix alone. The heterogeneity of the niche overlaps reappears only at next order, and there it acts not as a source but as a contamination, transmitting the mismatch of one pair to the correlation of another.

### E. In diverse communities, the centered mismatch dominates

In practice, there is little reason to expect the similarity in environmental responses to match the similarity in resource use closely enough to produce exact cancellation. ***β*** may deviate from zero. One might hope that this imbalance, small and unsystematic though it may be, still allows to infer niche overlap. In fact, as long as the *heterogeneity* of ***α***, i.e., the pair-to-pair variation in interaction strength, is considered, it does not. The reason is already visible in Eq. (4): the interaction matrix enters the correlations only through the propagator, not as a source, and to the leading order the propagator is built from the *homogeneous* part of ***α*** alone. The pairwise overlaps are thus a very weak signal, and recovering them would in any case require knowing ***β***, which is tied to no accessible observable.

One might then expect that in a large community even the dependence on the mismatch ***β*** should wash out: with each species buffeted by the fluctuations of all *S* − 1 others, the correlation of any particular pair should average away, of order 1*/S*. Again, the opposite is true. Solving Eq. (4) for a homogeneous ***α***, the legitimate leading-order limit, gives at large *S*

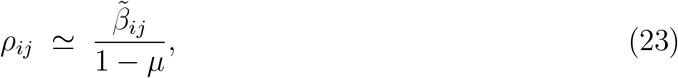

an *O*(1) quantity, where 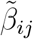 is the centered mismatch of Appendix E, independent of ***α***. The degeneracy does not close with diversity, it sharpens: in a large community *ρ*_*ij*_ becomes a clean readout of that pair’s residual mismatch, and of the pairwise residual overlaps *δα*_*ij*_ it retains nothing. This is demonstrated in Fig. 3.

**FIG. 3.**
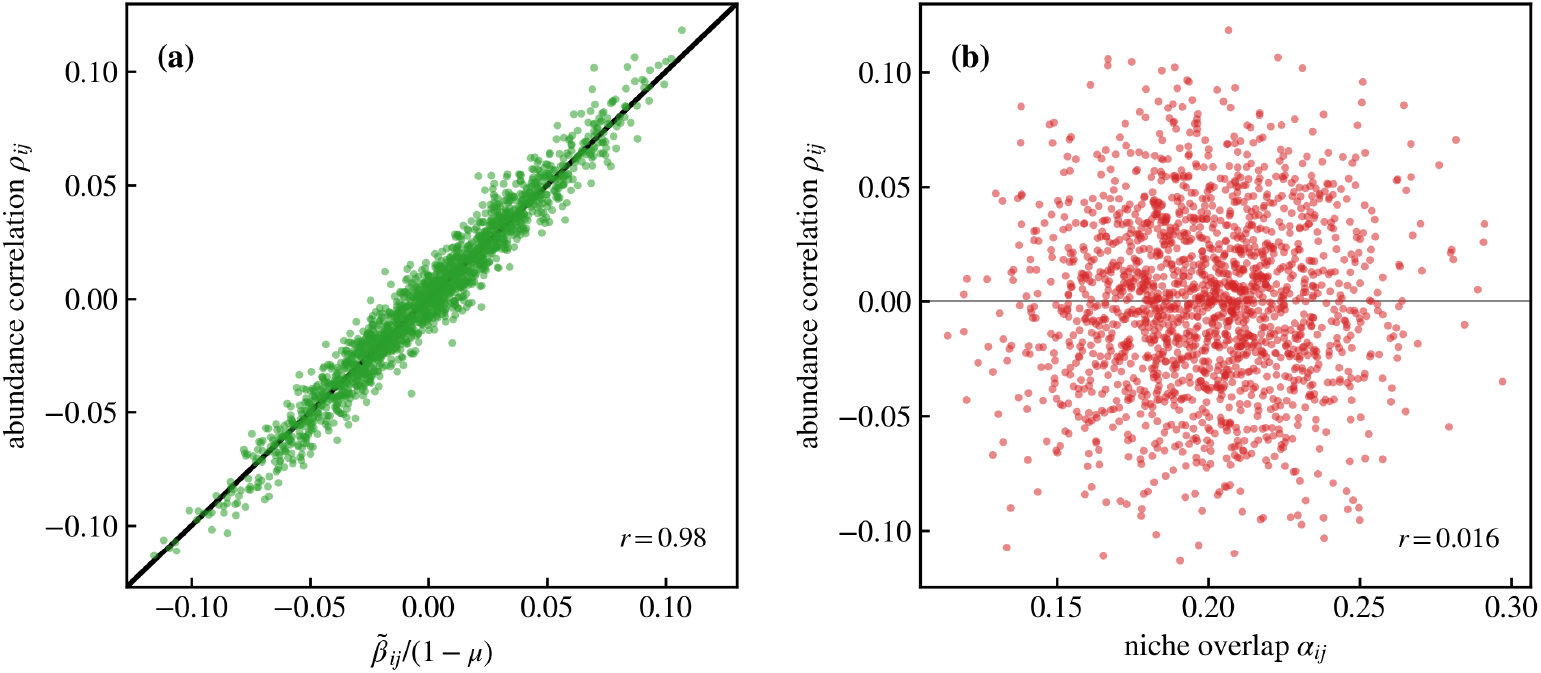
Abundance correlations report the mismatch, not the niche overlap. A community of *S* = 60 species with heterogeneous symmetric interactions, *α*_*ij*_ = *µ* + *δ*_*ij*_ and a response–competition mismatch *C*^(*η*)^ = ***α***_*S*_ + ***β***. The heterogeneity *δ*_*ij*_ = *δ*_*ji*_ and the mismatch *β*_*ij*_ = *β*_*ji*_ are drawn independently, each from N(0, *σ*^2^) with *σ* = 0.03, and *µ* = 0.2 . Each point is one of the 1770 species pairs, correlations are exact linear-response results, obtained by solving Eq. (4). **(a)** *ρ*_*ij*_ against the prediction of Eq. (4), the pair’s centered mismatch divided by the softness factor 1 − *µ*: the correlation of a pair is a direct readout of the centered mismatch of *that* pair. **(b)** The same *ρ*_*ij*_ against the niche overlap *α*_*ij*_, the quantity the measurement is supposed to report. There is no relation.

## V. RETARDATION: WHEN COMPETITION LAGS THE SHARED RESPONSE, CORRELATIONS MAY TRACK NICHE OVERLAP

In the last section we discovered that, at the biologically grounded point, *C*^(*η*)^ = ***α***_*S*_, the abundance correlations vanish identically. As explained, this reflects cancellation: the negative correlations due to competition cancel the positive correlations that reflect similar response to the environment.

The cancellation derived above relies on competition and shared environmental forcing entering the effective Lotka–Volterra dynamics on the same timescale. This raises a natural question: what happens when competitive feedback is mediated by resources with a finite response time?

This phenomenon cannot be examined within the Lotka–Volterra model, which contains no clock: the resources have been eliminated, and with them the delay. We therefore return to the consumer–resource dynamics of Sec. II, now made stochastic. Furthermore, we retain the resource timescale *τ*_*R*_ explicitly:

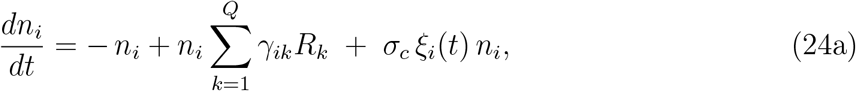

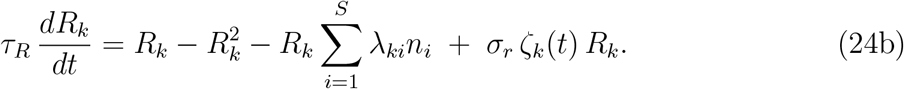

The environment now enters through *two* distinct channels, and the two are separately switchable. The consumer noise *ξ*_*i*_ acts on the growth rates directly, with

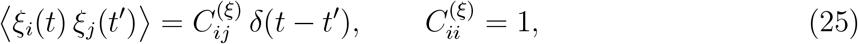

and represents mechanism (A) of Sec. IV: species of similar physiology respond similarly to temperature, humidity or pH, and *C*^(*ξ*)^ is where that similarity is encoded. The resource noise *ζ*_*k*_ acts on the supply,

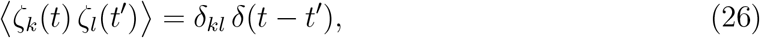

and represents mechanism (B): the resources fluctuate independently of one another, and the consumers inherit whatever correlation their shared reliance on them implies. Note that nothing is assumed about the consumers in this second channel, the covariance they experience is not imposed but *derived*. Setting *σ*_*r*_ = 0 isolates (A), setting *σ*_*c*_ = 0 isolates (B). It is this separation, impossible in a model with no resources in it, that the remainder of this paper exploits. In the deterministic limit, and for *τ*_*R*_ → 0, Eqs. (24) reduce to the Lotka–Volterra system of Sec. II with ***α*** = ΓΛ.

To test the effect of retardation we set *σ*_*r*_ = 0, so that the environment acts on the consumers alone. Imposing the grounded prescription that annihilated the signal one level of description higher,

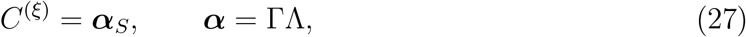

we find (Fig. 4a) that the correlations no longer vanish, on the contrary, they increase approximately linearly with the niche overlap, under the very assumption that, in the Lotka– Volterra description, forces *ρ*_*ij*_ = 0 identically. Hence, if competition is delayed relative to the shared response, and if the noise covariance tracks the interactions, as suggested by the plausibility argument (A) above, then abundance correlations may indeed serve as a good proxy for *α*_*ij*_, and may decay with phylogenetic distance (assumed to reflect niche overlap) in the manner reported by Sireci *et al*. [2].

**FIG. 4.**
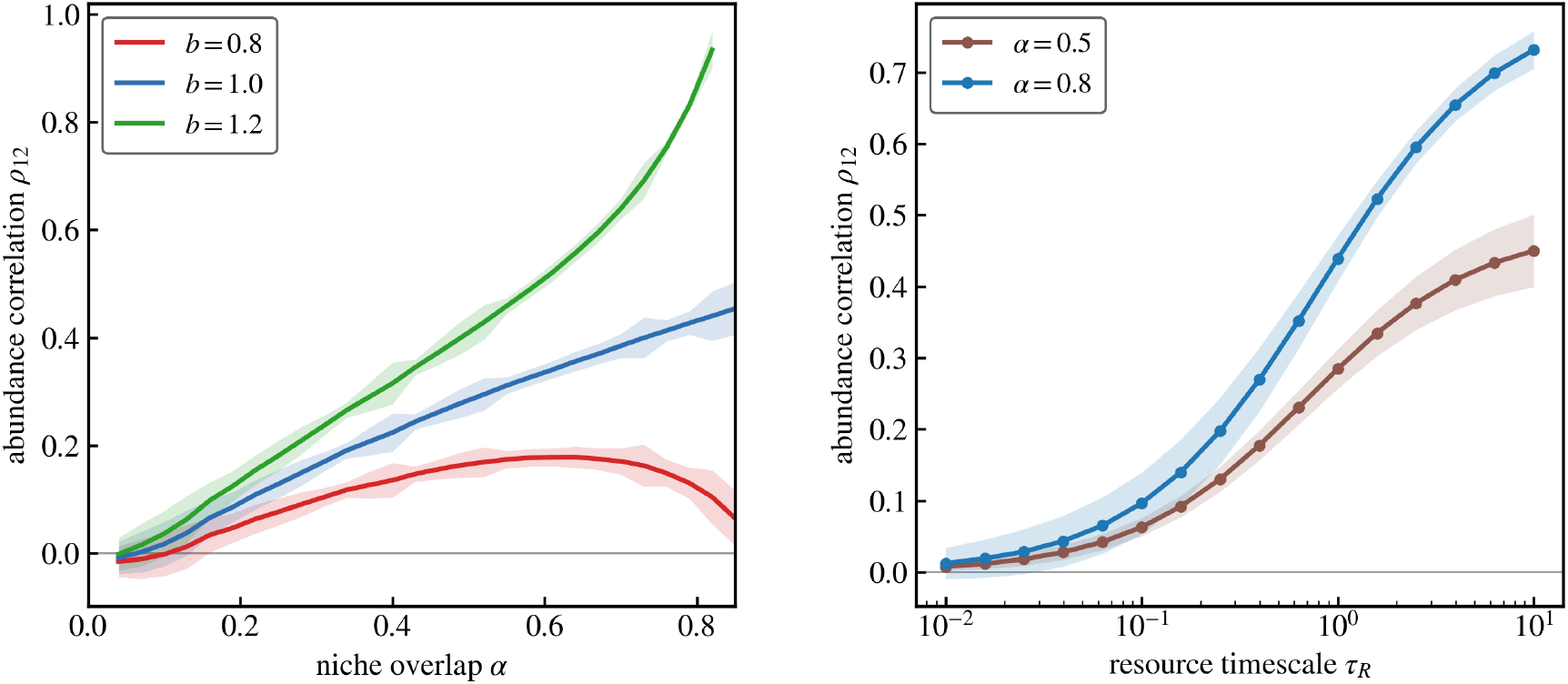
Retardation exposes the shared response, and removing it restores the cancellation. Abundance correlation between two consumers in the stochastic consumer–resource system, Eq. (24), with the environment acting on the consumers alone (*σ*_*r*_ = 0, *σ*_*c*_ = 0.05), *S* = 2 and *Q* = 3. Solid curves are means over 100 accepted (Γ, Λ) realizations that reproduce the same niche-overlap matrix, shaded bands are one standard deviation across those realizations. **(a)** Correlation against the niche overlap *α*, at resource timescale *τ*_*R*_ = 1, for a noise correlation 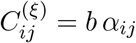. At the biologically grounded point *b* = 1, where, in the Lotka–Volterra description, *ρ*_*ij*_ = 0 identically for every overlap (Figure 2), the correlation is instead positive and grows steadily with *α*, because competition is transmitted through the resource pool and therefore lags the shared environmental response. The curves at *b*≠ 1 show that retardation outruns competition rather than overwhelming it: at *b* = 0.8 the correlation is positive at moderate overlap but bends back toward zero as *α* grows and competition reasserts itself. **(b)** The crucial role of retardation. Holding *b* = 1 and varying the resource timescale *τ*_*R*_ over three decades, at two values of the niche overlap. As *τ*_*R*_ → 0 the resources are slaved instantaneously to the consumers, the reduced dynamics is exactly the stochastic Lotka–Volterra system with ***β*** = 0, and the cancellation is restored, monotonically, and to zero. The correlation is therefore a property of the delay, not of the consumer–resource description as such.

Two reservations follow. First, the effect is genuinely one of retardation: removing the delay removes the correlations. As *τ*_*R*_ → 0 the resources are slaved instantaneously to the consumers, the reduced dynamics is precisely the stochastic Lotka–Volterra system with ***β*** = 0, and the cancellation is restored in that limit. Figure 4b shows this over three decades in *τ*_*R*_. Second, the vanishing of ***β*** relies on the exact fulfillment of the biological argument (A) of Sec. IV. Once it does not hold exactly, the one-to-one relationship between correlations and interactions breaks down (Fig. 4a) and can even become non-monotonic. The interpretation therefore requires two pieces of information that the equal-time correlation matrix does not itself provide: whether competitive feedback is appreciably retarded, and how closely the covariance of direct environmental forcing follows the niche-overlap matrix.

## VI. RESOURCE-MEDIATED NOISE: A SIGNAL SET BY THE YIELD–DEPLETION MISALIGNMENT, NOT BY *α*

Let the resources fluctuate, independently of one another, and let the consumers feel the environment only through what they eat: *σ*_*c*_ = 0, *σ*_*r*_ *>* 0 in Eq. (24). This is mechanism (B) of Sec. IV in its bare form. Nothing is imposed on the consumers and the covariance they experience is not stipulated but *derived*. The question is what the resulting abundance correlations tell us.

Eliminating the resources adiabatically returns, exactly, a stochastic Lotka–Volterra system (Appendix F). Its interaction matrix is the usual one, but the environmental noise it inherits is not:

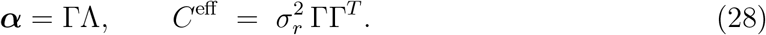

Competition is built from yield and depletion, the inherited forcing from yield alone. The two are different combinations of the same traits, and their difference plays the role of the mismatch of Sec. IV,

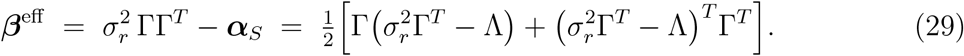

The correlations are therefore governed not by the niche overlap but by the misalignment between what a consumer takes from a resource and what it gets out of it.

The two matrices in Eq. (4) differ in a second way, less obvious than the first. The interaction matrix is normalized, *α*_*ii*_ = 1, whereas the inherited covariance is not: its diagonal entries 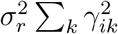 differ from one species to the next. This is not merely a matter of overall scale. Our normalization fixes the *sum* of each yield profile, ∑_*k*_ *γ*_*ik*_ = 2 (Sec. II B), but the inherited amplitude is set by the sum of squares. Under a fixed row sum the latter is largest for a concentrated profile (yield specialist) and smallest for an even one (yield generalist). Two species with identical niche overlap, and hence identical ***α***, can differ several-fold in this amplitude, a distinction the Lotka–Volterra description, which normalizes it away, cannot see. For clarity of presentation, we neglect this difference in the discussion here. The exact expression for ***β***^eff^ in the case of a non-uniform noise diagonal is given in Appendix G.

Two consequences follow.

First, when the yield is proportional to the depletion, Λ = *c* Γ^*T*^, the assumption on which most consumer–resource theory rests, there is no signal [2, 15, 16]. In that case ***α*** = *c* ΓΓ^*T*^ and 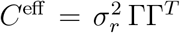 are proportional to the *same* matrix. The normalization *α*_*ii*_ = 1 then forces 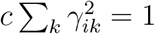 for every *i*: the diagonal of ΓΓ^*T*^ is uniform, and with it the inherited noise amplitudes, which are common to all species. What is left is a single scalar, 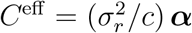, and an overall factor cannot affect a correlation: every interspecific correlation thus vanishes. Resource fluctuations, however strong, produce no abundance correlations at all, the shared environmental response and the competition have become the same object, and cancel as they did in Sec. IV.

Second, when yield and depletion are misaligned, there is a signal, but not about ***α***. Figure 5 demonstrates that while making the contrast with consumer noise directly. Under consumer noise, the correlations collapse onto a single increasing function of the niche overlap. Under resource-mediated noise, that collapse is destroyed: communities with *identical* niche overlap yield correlations of either sign, ordered by the yield–depletion mismatch *D* of Ref. [16]. This is a per-pair quantity, like ***β***, but of different origin: ***β*** is the source term of the Lyapunov equation that fixes the covariance, and hence the correlations, whereas *D* is a guess at the correlations themselves, read off from an exactly solvable two-species example. In the exactly solvable case of Appendix G the two change sign together, so that either serves equally well as a qualitative predictor.

**FIG. 5.**
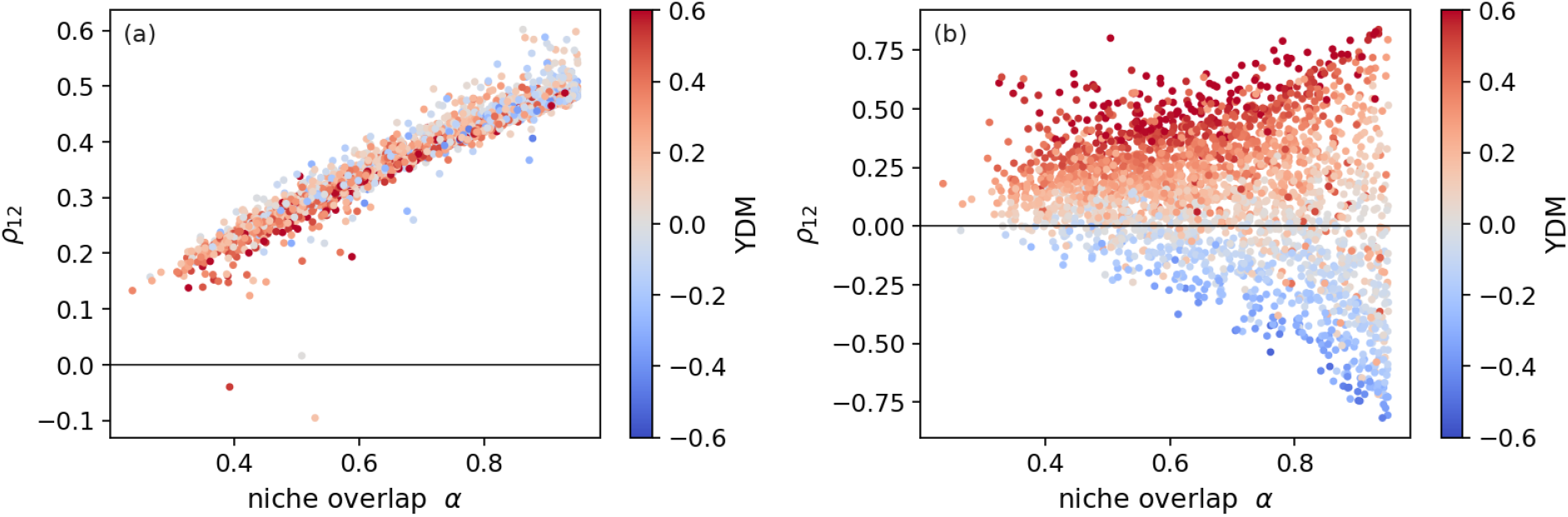
The same niche overlap, correlations of either sign. In both panels each point is one accepted (Γ, Λ) realization of a two-consumer, three-resource community (*τ*_*R*_ = 1), the color encodes the yield–depletion mismatch *D* of Ref. [16]. **(a)** Direct consumer noise (*σ*_*r*_ = 0, *σ*_*c*_ = 0.05, *C*^(*ξ*)^ = ***α***_*S*_). The points collapse onto a single increasing curve, and the colour is unstructured: the correlation is a function of the niche overlap, and the mismatch is irrelevant. This is the retardation regime of Sec. V. **(b)** Resource-mediated noise (*σ*_*r*_ = 0.05, *σ*_*c*_ = 0). The collapse is destroyed. At any fixed *α* the correlation ranges over both signs, and the colour separates the cloud almost completely: what orders the points is the yield–depletion mismatch, not the overlap. Communities that compete identically, same ***α***, same fixed point, are strongly correlated, uncorrelated, or strongly anticorrelated, according to a property of Γ and Λ that ***α*** does not contain.

Resource-mediated noise, then, is informative, but not about the niche overlap. Rather, it informs us about the internal economy of resource use. Two communities with the same niche overlap structure may still be positively correlated, uncorrelated, or negatively correlated, according to a property of the yield–depletion relation.

## VII. DISCUSSION

Many theoretical studies have examined stochastic community dynamics, yet the relationship between niche overlap and the correlation structure of environmental fluctuations has not been analyzed systematically. Broadly speaking, there are two natural ways to model environmental stochasticity subject to these relationships. One is to model fluctuations at the level of the resources, allowing the induced correlations between species to emerge from the dynamics themselves. The other is to prescribe externally a correlation structure that reflects, at least approximately, the similarity in species’ resource use.

Here we have shown that under this natural benchmark the equal-time abundance correlations vanish exactly, and that even small departures from it do not recover information about the interaction matrix. Equal-time correlations become informative about interactions only when environmental fluctuations act directly on the consumers with an appropriate temporal delay. When fluctuations act on the resources, they instead report the mismatch between resource yield and depletion. In practice, however, the source of environmental variability, the presence and duration of delays, and related details are often unknown. Without such information, equal-time abundance correlations provide only limited insight into the underlying ecological interactions.

### A. Relation to dynamical inference methods

Through this paper we considered only equal-time abundance correlation. It is the most accessible of the fluctuation statistics: it requires only that abundances be recorded, and being an average it is comparatively robust to measurement error. Two other families of inference read more from the same fluctuations, and it is worth placing our result beside them.

The first estimates the interaction matrix from the *rate of change* of the abundances, fitting the growth rate to the abundance, or equivalently taking a short-lag derivative of the covariance [34–37]. In the white-noise limit this is exact: writing ∑(*τ*) = ⟨*δn*(*t*+*τ*) *δn*(*t*)^*T*^ ⟩ = *eJ*^*τ*^ ∑(0) for *τ >* 0, the one-sided derivative at *τ* = 0^+^ gives the drift *J* = −*N* ***α***. Therefore, 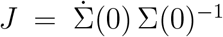. The degeneracy we have described is broken, because the derivative supplies the second moment 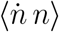 that the correlation alone omits.

But the exactness is fragile, in three independent ways. When the environmental forcing carries temporal structure, as ecological forcing generally does, the cross-term ⟨*η n*⟩ no longer vanishes and the estimate is biased. Because it rests on a numerical derivative, it amplifies measurement error, inflating the high-frequency content where noise dominates. Even with white noise and perfect data, the derivative must be taken at a finite sampling interval, adequate only for modes whose relaxation time it resolves: a real community carries a *spread* of relaxation timescales, and a single finite sampling interval may fail to resolve them unless the temporal resolution and the record length together span the full dynamical range. What is a virtue of the equal-time correlation, that it is a stationary quantity, integrating over time rather than differentiating, and indifferent to the sampling rate, is precisely what these methods give up.

A second family, developed by Chen *et al*. [4], reads the fluctuations not through the equal-time correlation but through their full frequency content, using the cross-spectrum and the coherence. This is a genuinely different probe, aimed at a different target: not the interaction matrix ***α*** = ΓΛ, but the structure of shared resource use ΓΓ^*T*^, which guild a species belongs to, rather than how strongly it competes. In our decomposition that target is the response channel, the very object the correlations do carry, and the two coincide only near the point where yield and depletion are proportional. Their method relies on the environment being temporally structured, the coherence between two species is the signature of their common, colored, resource-mediated drive.

The three approaches are therefore complementary rather than competing: they read different moments of the same fluctuations, and they recover different things, the mismatch, the interactions under white forcing, and the resource-sharing structure, respectively. But all three differ only in *which* statistic they read. A separate question is when the equal-time correlation, the humblest of them, nonetheless becomes informative, and it does so not by reading more, but by importing an independent ordering of the pairs.

### B. Mechanistic interpretation with external information

The single case in which the correlation matrix speaks for itself is the retardation regime, and even there the resource timescale that licenses the reading cannot be read off the abundances. One might expect the resource dynamics to leave a second, faster branch in the consumer autocorrelation, but it does not: the resources are slaved to the consumers and their fluctuations are an echo rather than a source, so the fast modes carry negligible weight. The resource timescale does shift the slow relaxation rates, but it shifts them together with ***α***, and the two cannot be separated without knowing one of them in advance.

The correlations may become informative, though, as soon as one supplies an independent ordering of the pairs by niche overlap. Suppose only that relatedness is a proxy for niche overlap, so that *α*_*ij*_ decreases with the phylogenetic or genetic distance between species *i* and *j*. This is an assumption about the *interactions*, not about the noise. Granted it, the correlation matrix becomes a readout of the machinery that produced it.

For example, if the measured *ρ*_*ij*_ fall off with phylogenetic distance, as reported by Sireci *et al*. [2], it implies, first, that the community does not sit at the grounded point, where Eq. (4) forces every correlation to vanish. Moreover, we discovered that resource-mediated forcing produces correlations organized by the yield–depletion mismatch rather than by the overlap (Fig. 5b), whereas direct forcing on the consumers, with competition retarded by the resource pool, produces exactly the observed ordering (Fig. 4a). A correlation that tracks relatedness therefore says: the environment acts on the consumers, and the resources are slow.

As another example, suppose instead that the environmental driver is known independently to enter through the resources, as in a chemostat, or wherever the fluctuating quantity is the nutrient influx. Then Eq. (4) applies, and correlations are possible if and only if depletion is not proportional to yield. An observed *ρ*_*ij*_ ≠ 0 establishes, therefore, that what a species removes from a resource is not proportional to what it gains from it. The sign and magnitude then quantify the misalignment. This is a mechanistic statement about resource use, obtained from an abundance time series.

A null result is not empty either. Under the biologically grounded prescription it is the prediction of two quite different situations, fast resources with consumer forcing, or resource forcing with yield proportional to depletion, and in both the community is telling us that it sits at or near the point where the environmental responses of two species resemble one another exactly as much as their resource use does.

Once the overlap is identified (e.g., from genetic distance as a proxy) the two mechanisms, consumer forcing and resource forcing, can be separated from the data. Under retardation, pairs of equal relatedness are equally correlated and, more to the point, they are correlated with the same sign: the correlation points collapse onto a single increasing function of *α*_*ij*_ (Fig. 5a). Under resource-mediated forcing the collapse is destroyed (Fig. 5b), and at any fixed overlap the correlations range over both signs, ordered by a quantity the overlap does not contain.

The statistic to report is therefore not the trend of *ρ* against relatedness, nor even the width of the scatter about that trend, which can be comparable in the two cases. It is whether pairs of equal relatedness are correlated in opposite directions. Both readings are available from an ordinary abundance time series, once the pairs have been sorted by relatedness.

What abundance correlations report, then, is not the interaction matrix but the route by which the environment reaches the community, and the internal economy of resource use along that route. This is less than has been claimed for them, and more than we had any right to expect.

## Acknowledgments

N.M.S acknowledges support from the Israel Ministry of Science (Italy-Israel cooperation, grant no. 7578) and of the Israel Science Foundation (grant no. 2435/24).

## Appendix A Generating Γ and Λ

Everything in Secs. V and VI rests on being able to produce many pairs (Γ, Λ) that yield the *same* niche-overlap matrix ***α*** = ΓΛ while differing in the yield–depletion relation. This appendix specifies how that is done.

We adopted an approach in which both matrices were determined together, by constrained optimization, and the nonnegativity is imposed as a bound rather than checked afterwards.

Given a target interaction matrix ***α***, with *α*_*ii*_ = 1 and a prescribed mean overlap, we minimize

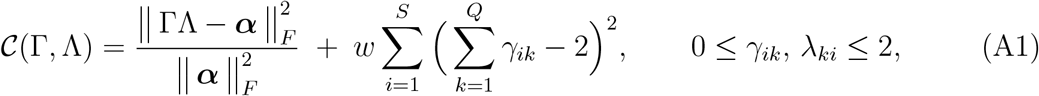

by sequential quadratic programming, from a starting point drawn uniformly on [1, 2]. We take *w* = 0.1.

The first term asks that the pair reproduce the required niche overlap. The second enforces the normalization of Sec. II B: rows of Γ summing to 2 give *r*_*i*_ = ∑_*k*_ *γ*_*ik*_ − 1 = 1, so that all species share the same intrinsic growth rate.

Different random starting points converge to different (Γ, Λ) that reproduce the *same* ***α***. This is the degeneracy the paper is about, and it is what supplies the spread of yield– depletion mismatch at fixed niche overlap in Figs. 4 and 5.

### 1. What the procedure delivers

Because Eq. (A1) is minimized rather than solved, the two requirements compete, and neither is met exactly. Both must therefore be reported rather than assumed. Table A1 gives them, over 40 independent runs for each parameter set.

### 2. Rejection criteria

A pair (Γ, Λ) that reproduces ***α*** need not describe a viable community. We discard it unless

i. *λ*_*ki*_ ≥ 0 for all *k, i* (guaranteed by the box constraints),
ii. the coexistence fixed point is positive, 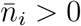 for all *i*,
iii. the quasi-steady resource abundances are positive, 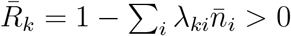 for all *k*,
iv. that fixed point is linearly stable.

**TABLE A1.** : Accuracy of the sampler, median over 40 runs. The worst case over all runs and all parameter sets is 2 × 10^*−*6^ in each column. The reconstructed interaction matrix agrees with the target, the rows of Γ sum to 2 so that *r*_*i*_ = 1, and the diagonal of ΓΛ is unity, all to within numerical tolerance. The residual is small enough that the intrinsic growth rates may be treated as uniform: the spurious correlations generated by growth-rate heterogeneity scale as its square, and are here of order 10^*−*12^.

| $S$ | $Q$ | $\alpha$ | $\frac{\ \Gamma\Lambda - \alpha\ _F}{\ \alpha\ _F}$ | $\max_i \vert \sum_k \gamma_{ik} - 2\vert $ | $\text{diag}(\Gamma\Lambda)$ |
| --- | --- | --- | --- | --- | --- |
| 2 | 3 | 0.6 | $1.4 \times 10^{-7}$ | $1.7 \times 10^{-7}$ | 1.0000 |
| 2 | 3 | 0.4 | $1.2 \times 10^{-7}$ | $3.5 \times 10^{-7}$ | 1.0000 |
| 2 | 3 | 0.8 | $1.2 \times 10^{-7}$ | $2.5 \times 10^{-7}$ | 1.0000 |
| 5 | 8 | 0.3 | $5.5 \times 10^{-7}$ | $4.7 \times 10^{-7}$ | 1.0000 |

Acceptance is high: 40*/*40 for *S* = 2, *Q* = 3 at mean overlap 0.4 and 0.6, 39*/*40 at 0.8, and 40*/*40 for *S* = 5, *Q* = 8 at mean overlap 0.3. The rejections occur at strong overlap, where 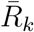 approaches zero.

## APPENDIX B Derivation of the reduced Lyapunov equation

This appendix derives Eq. (4), on which the whole of Sec. IV rests, and establishes the uniqueness of its solution. No symmetry of ***α*** is assumed at any point.

### 1. The reduction

We start from the Lyapunov equation (11) with the drift and diffusion matrices of Eq. (4),

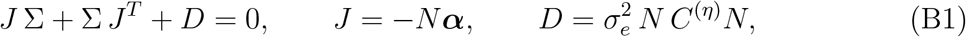

and substitute the ansatz

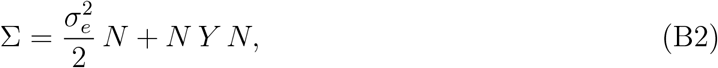

with *Y* symmetric. Expanding,

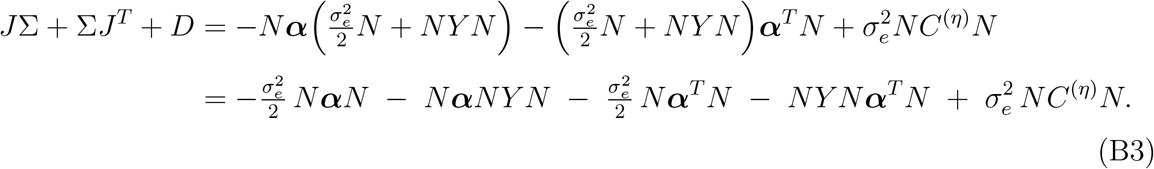

Every term carries a factor *N* on the left and a factor *N* on the right. Since *N* is diagonal with strictly positive entries at a coexistence fixed point, it is invertible, and we may multiply through by *N* ^*−*1^ from both sides. Setting the result to zero and rearranging,

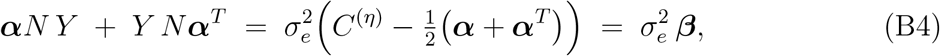

which is Eq. (4).

Two features of this calculation are worth isolating.

- **The symmetrization is forced, not chosen**. The combination 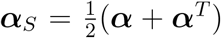 arises from a single source: the first term of the ansatz, 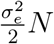, contributes 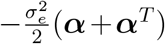 after the factors of *N* are stripped, because the Lyapunov equation contains both *J* Σ and its transpose. It appears nowhere else. The symmetric part of the interaction matrix is therefore not an assumption imposed for convenience, and not a definition of niche overlap adopted in advance: it is the only part of ***α*** that the equation permits a noise covariance to be compared with. A covariance matrix is symmetric, and the antisymmetric part of ***α*** has nothing symmetric to be measured against.
- **The interaction matrix is not a source**. In Eq. (B4) the matrix ***α*** appears twice on the left, acting on *Y*, and once on the right, inside ***β***. Its appearance on the left is that of an operator: it determines how a given source is transmitted through the community. Its appearance on the right is only as the reference against which the noise is measured. There is no term in which ***α*** drives the correlations by itself.

### 2. Existence and uniqueness

Write *K* = ***α****N*, so that Eq. (B4) reads

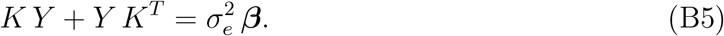

This is a Sylvester equation. The linearization presupposes a stable coexistence fixed point, and stability places every eigenvalue of *J* = −*N* ***α*** in the left half-plane, so every eigenvalue of *K* = ***α****N*, which shares the spectrum of *N* ***α***, has positive real part. No two can then sum to zero, and the solution is therefore unique for any ***β***, symmetric or not. If ***β*** = 0 then *Y* = 0 is that unique solution, so by Eq. (B2) the abundance covariance is 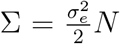, diagonal, and every interspecific correlation vanishes identically. The cancellation therefore holds around any stable fixed point, whatever the asymmetry of ***α***. Asymmetry bears only on whether the matched noise *C*^(*η*)^ = ***α***_*S*_ is itself admissible, which requires ***α***_*S*_ to be positive semidefinite (Appendix C 5).

### 3. Why the heterogeneity of *α* does not propagate

Decompose the interaction matrix into a homogeneous part and a perturbation, 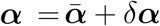, with 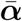 having unit diagonal and all off-diagonal entries equal to the mean overlap *µ*. The fixed point shifts with it,

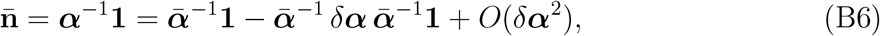

so that 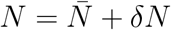 with *δN* = *O*(*δ****α***).

Now observe that the source term of Eq. (B5) is *O*(***β***), and hence so is *Y* . Both *δ****α*** and *δN* enter the equation only through the operator on the left, that is, only multiplied by *Y* . Their contribution is therefore *O*(*δ****αβ***), and to leading order in the mismatch

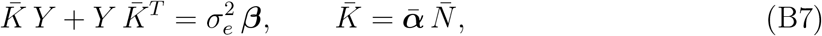

in which the propagator is built from the *homogeneous* interaction matrix alone. The pairwise heterogeneity of the niche overlaps, which is the quantity the inference is trying to recover, is absent from the leading-order answer. It reappears at second order, and there it does not act as a source but as a contamination: it transmits the mismatch of one pair to the correlation of another along indirect paths *i* → *k* → *j*.

### 4. Confirmation by direct simulation

Equation (B4) is exact within the linearized weak-noise description. Since the underlying dynamics, Eq. (4), is nonlinear and the noise multiplicative, we confirmed it against direct Stratonovich simulation of the full stochastic process for two species. The estimated correlations converge to the values predicted by Eq. (4). In fact, at the matched point the cancellation is exact and non-perturbative in the noise strength, as we prove in Appendix C.

## Appendix C Exact factorization at the matched point

The cancellation of interspecific correlations at the matched point *C*^(*η*)^ = ***α***_*S*_ [Eq. (4)] was obtained above from the linearized Lyapunov equation, that is, to leading order in the noise strength. At the matched point the result is in fact exact and non-perturbative: the full nonlinear stationary distribution factorizes into independent single-species Gamma laws, for any noise strength and for arbitrary, possibly non-reciprocal, interactions. We give the derivation here.

In what follows we use the general stochastic Lotka–Volterra system, with heterogeneous growth rates *r*_*i*_ and an arbitrary interaction matrix ***α***,

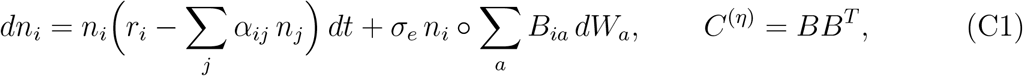

in the Stratonovich convention (denoted º), and assume a positive coexistence fixed point, 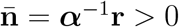, so that

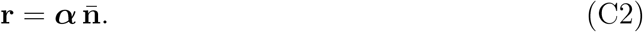

Eq. (4) is the special case *r*_*i*_ = 1. We develop the argument for the standard bilinear interaction *α*_*ij*_*n*_*j*_ and a common noise amplitude. A more general interaction term, and a species-dependent amplitude, are treated at the end of the appendix.

In the logarithmic variables *x*_*i*_ = log *n*_*i*_ the Stratonovich chain rule holds without a drift correction, and the multiplicative noise becomes additive with a constant coefficient,

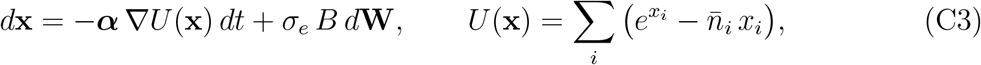

with constant diffusion 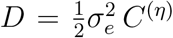. Importantly, the potential *U* is *separable*, a sum of single-species terms, because the logistic nonlinearity 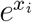 is diagonal in the site basis, and the interaction ***α*** enters only as a constant mobility, not through *U* .

### 1. Symmetric interactions

Take ***α*** = ***α***_*S*_ and the matched noise *C*^(*η*)^ = ***α***_*S*_, so that 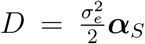. Mobility and diffusion then satisfy an Einstein relation, *D* ∝ ***α***, and the Boltzmann distribution

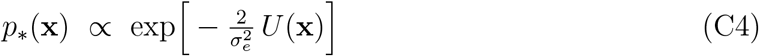

is stationary: with 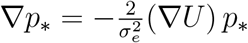 the probability current

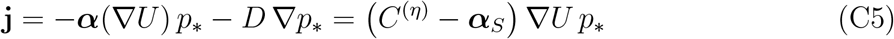

vanishes identically. More generally, for an arbitrary noise covariance the current is **j** = *C*^(*η*)^ − ***α*** (∇*U*) *p*_∗_, and the symmetric part *C*^(*η*)^ − ***α***_*S*_ controls whether the current vanishes (detailed balance). The antisymmetric part, minus ***α***_*A*_, persists even at the matched point, and this case is considered below.

Because *U* is separable, *p*_∗_ factorizes, and in the original variables

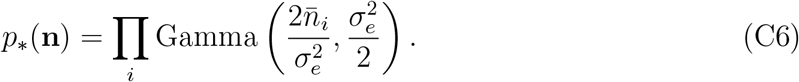

so each species is an independent Gamma variable with 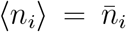 and 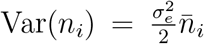. Independence gives ⟨*n*_*i*_*n*_*j*_⟩ = ⟨*n*_*i*_⟩⟨*n*_*j*_⟩ and hence

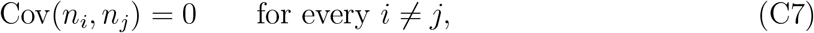

exactly and for arbitrary *σ*_*e*_.

The factorization is in fact stronger than the vanishing of the covariance, since all mixed cumulants and the equal-time mutual information vanish as well. The matched noise exists provided ***α***_*S*_ ⪰ 0 (the symbol ⪰ stands for positive semi-definite), which holds automatically for a symmetric ***α*** with a stable positive fixed point.

A factorized Gamma stationary distribution under a detailed-balance relation between the interaction and environmental-noise matrices was previously derived by Camacho-Mateu et al. [15]. In what follows we extend the factorized solution to nonreciprocal interactions, for which the stationary state may support a nonzero, but divergence-free, probability current.

### 2. Non-reciprocal interactions

Now let ***α*** = ***α***_*S*_ + ***α***_*A*_ with ***α***_*A*_ ≠ 0. As long as ***α***_*S*_ is positive definite, i.e., as long as a community with ***α***_*S*_ alone is feasible and stable (although with a different set of equilibrium abundances) it can still define a correlation matrix, so one may choose *C*^(*η*)^ = ***α***_*S*_. Although the covariance can match only ***α***_*S*_, the *same p*_∗_ of Eq. (C4) remains the stationary probability distribution function.

The only difference is that, once the interaction matrix is asymmetric, Eq. (C4) no longer satisfies detailed balance. Instead, it now carries a nonzero current,

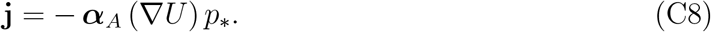

This is the ***α***_*A*_ term identified in the general decomposition above: the symmetric part of the current vanishes because *C*^(*η*)^ = ***α***_*S*_, leaving only the antisymmetric residual.

Despite carrying a current, the probability distribution (C4) is still time independent, because this current is divergence-free. Writing *j*_*i*_ = − ∑ _*k*_ (*α*_*A*_)_*ik*_ (*∂*_*k*_*U*) *p*_∗_,

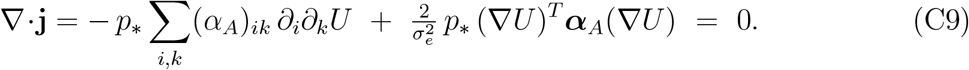

Both terms in this equation vanish by the antisymmetry of ***α***_*A*_ alone: the first is the contraction of the antisymmetric ***α***_*A*_ with the symmetric Hessian *∂*_*i*_*∂*_*k*_*U*, the second the quadratic form (∇*U*)^*T*^ ***α***_*A*_(∇*U*). Neither step uses the separability of *U*, so the stationarity of *p*_∗_ holds for any potential. Hence *p*_∗_ remains stationary, and the factorization (C6), with Cov(*n*_*i*_, *n*_*j*_) = 0, holds verbatim for the full, non-reciprocal ***α***.

As mentioned, detailed balance is now broken: the steady state supports a persistent rotational probability current and is irreversible. The interaction is thus invisible to the equal-time distribution while remaining fully present in the dynamics, in lagged correlations and in the entropy production of the abundance process. Separability enters only in the last step of the symmetric case, in reading off the factorized form. The stationarity of *p*_∗_ does not require it.

### 3. Generalizations

- *General interaction*. The bilinear form *α*_*ij*_*n*_*j*_ may be replaced by *α*_*ij*_ *ψ*_*j*_(*n*_*j*_) with an arbitrary species-specific monotone *ψ*_*j*_, one per species (a Holling saturation *ψ*_*i*_(*n*) = *n/*(1 + *k*_*i*_*n*) is one example). Each *ψ*_*j*_ is absorbed into a separate coordinate *V*_*j*_ of the potential *U* = ∑ _*i*_ *V*_*i*_(*x*_*i*_), which stays separable, so the factorization persists, now with non-Gamma marginals set by *ψ*_*i*_. The derivation uses only that the nonlinearity be *elementwise*: species *j* acts through a function of *n*_*j*_ alone. An aggregate nonlinearity, *g* (∑_*j*_ *α*_*ij*_*n*_*j*_), generally destroys separability. Whether its stationary law remains approximately factorized is a separate question.
- *General noise*. The matched noise of Eq. (4) has a unit diagonal, which presumes a common forcing amplitude across species. What if different species suffer from noises of different amplitude, i.e., when the covariance matrix is not unit diagonal?

Let us consider the case in which the diagonal terms of the covariance matrix of the noise are *H* = diag(*h*_1_, …, *h*_*S*_), instead of 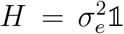. We would like to show that here also one can find a noise covariance matrix that supports a separable stationary distribution function, hence all zero-time interspecific correlations vanish.

To achieve that we choose a noise covariance

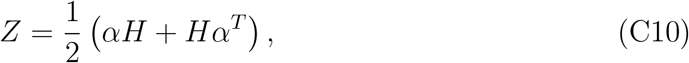

which is symmetric by construction. Note that the drift in log variables 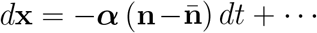 is unchanged: it is determined by the interaction matrix ***α*** and the fixed point 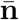, and does not depend on *H*.

We then define

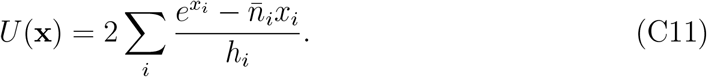

Hence,

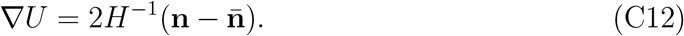

Substituting these expressions into the probability-current equation,

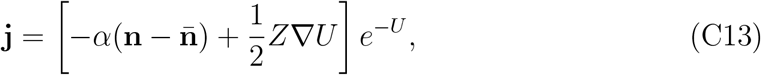

we obtain

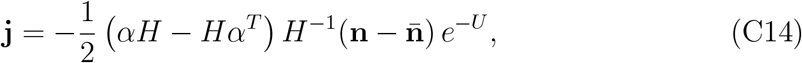

which is divergence-free:

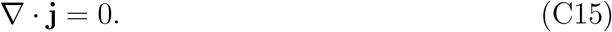

The construction so far only requires *Z* to be symmetric. For it to be a genuine noise covariance one also needs admissibility, the existence of a real *B* with 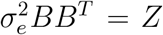. This is more restrictive than ***α***_*S*_ ⪰ 0: it requires the *H*-weighted symmetric part 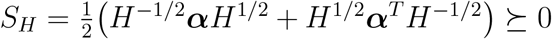. However, when this condition is satisfied, once again the stationary probability density factorizes into a product of Gamma distributions:

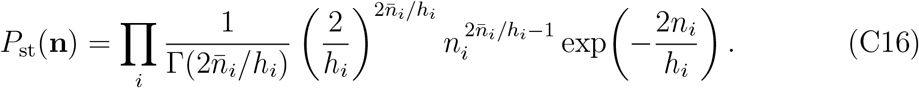

Equivalently,

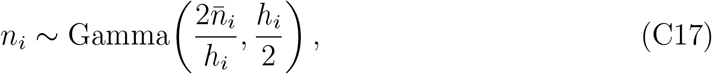

where the second argument is the scale parameter.

This case, different levels of stochasticity for different species, manifests itself when the forcing is resource-mediated, as in (Sec. VI). 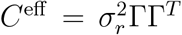 has a non-uniform diagonal, *H* = diag(*C*^eff^), and its exact-cancellation condition is Eq. (C10).

- *Itô convention*. The derivation is in the Stratonovich convention, natural if the white noise is the short-correlation limit of an external forcing. In the Itô convention the same factorization survives, with the Gamma means shifted by the usual noise-induced drift, 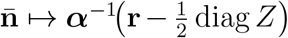. The difference is a shift of the coexistence point, not a change in the result. This shift is the multi-species counterpart of the mean 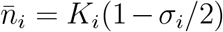 of the single-species logistic model with carrying capacity *K*.

### 4. The Gamma distribution

The product-form stationary distribution is thus an alternative route to the Gamma abundance laws familiar from neutral models and from non-interacting stochastic-logistic models [5, 13]: there the independent Gamma marginals arise because the species do not interact, or interact only through a mean field, whereas here they arise for an arbitrary interaction matrix, provided the environmental noise is matched to it. The Gamma abundance distributions and the vanishing equal-time correlations are all properties of the noise structure, not of the interactions. The independence of the equal-time marginals in particular is not a signature of the absence of interactions. It is a signature of the fluctuation–dissipation balance *C*^(*η*)^ = ***α***_*S*_ between the noise and the competition. The interactions are not absent, they are hidden: present in the dynamics, in the lagged correlations, and, for non-reciprocal ***α***, in the entropy production of the irreversible steady state, yet invisible to the equal-time distribution. One may conjecture that in the close neighborhood of the matched point the single-species abundances remain approximately Gamma distributed while the interspecific correlations already turn on, positive or negative according to the sign of the mismatch *β*, so that standard single-species distributions and nontrivial pairwise correlations coexist, as observed empirically and reproduced by the interaction-based model of Camacho-Mateu *et al*. [5].

### 5. Admissibility of the matched noise

Throughout this paper we assume that the noise covariance prescribed at the matched point is admissible, namely, that it is symmetric and positive semidefinite and can therefore be realized as the covariance of a multivariate noise process. We now make this condition explicit.

Consider first the normalization 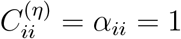. The matched noise correlation matrix is then

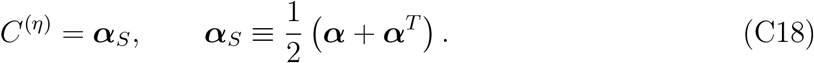

Consequently, the matched noise is admissible if and only if ***α***_*S*_ ⪰ 0.

For symmetric interactions, admissibility follows automatically from stable coexistence. Let 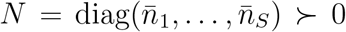, where the symbol ≻ implies positive definite, so that the Jacobian at the coexistence fixed point is *J* = −*N* ***α***. If the fixed point is linearly stable, all eigenvalues of *N* ***α*** are positive. Since ***α*** is symmetric, *N* ***α*** is similar to the symmetric matrix

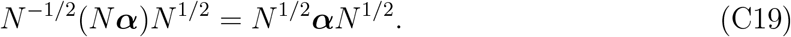

The latter therefore has only positive eigenvalues and is positive definite. Moreover, *N* ^1*/*2^***α****N* ^1*/*2^ is related to ***α*** by a congruence transformation. Sylvester’s law of inertia implies that a transformation of this type keeps the number of positive, negative and zero eigenvalues, therefore ***α*** ≻ 0. Thus, conditional on feasibility, stable coexistence for a symmetric interaction matrix implies that *C*^(*η*)^ ≻ 0, and the matched noise is admissible. Conversely, ***α*** ≻ 0 implies stability of the positive fixed point. Hence, for symmetric interactions and conditional on feasibility,

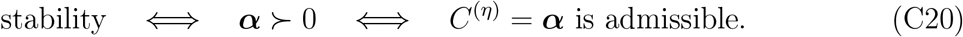

For asymmetric interactions this implication no longer holds. Stability of *J* = −*N* ***α*** does not, in general, imply ***α***_*S*_ ⪰ 0. With common forcing amplitudes, admissibility of the matched noise therefore requires the additional condition ***α***_*S*_ ⪰ 0. A sufficient condition is that the symmetric interaction matrix ***α***_*S*_, considered as a dynamical system in its own right, admit a stable positive fixed point.

More generally, let *H* = diag(*h*_1_, …, *h*_*S*_) ≻ 0 describe species-dependent forcing amplitudes. The covariance required at the matched point is

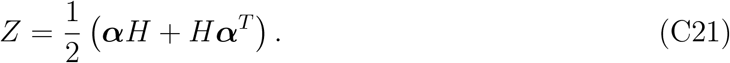

This construction defines a legitimate noise covariance only when *Z* ⪰ 0. Since *H* is positive diagonal, this condition is equivalently

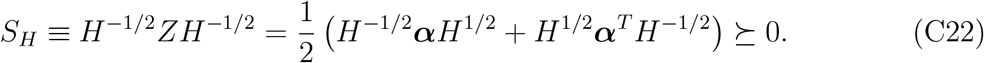

Thus, except in the symmetric common-amplitude case, dynamical stability and admissibility of the matched noise are distinct requirements.

## Appendix D Non-identifiability: sampling and admissibility

This appendix specifies the construction behind Fig. 1, and reports what it delivers. The algebra of Eq. (4) is exact and requires no verification. What does require verification is the one thing it does not guarantee: that the noise covariance it returns is a legitimate one, positive semidefinite and, once the free variances are fixed, of unit diagonal.

### 1. The construction

Given a candidate interaction matrix ***α*** and an observed covariance Σ^∗^, every subsequent step is a matrix operation and nothing is fitted:

i. the candidate fixes its own equilibrium, 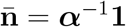, hence 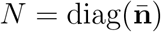 and *J* = −*N* ***α***,
ii. the diffusion follows from the Lyapunov equation, *D* = −(*J* Σ^∗^ + Σ^∗^*J*^*T*^),
iii. the noise covariance follows from the diffusion, 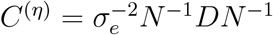.

By construction the candidate then reproduces Σ^∗^ exactly. This is not a numerical claim and there is nothing to check about it.

### 2. What must be checked

Steps (i)–(iii) return a symmetric matrix. They do not return a *covariance* matrix. For *C*^(*η*)^ to be admissible it must be positive semidefinite, and for it to be a correlation structure in the sense of Sec. III it must have a unit diagonal. Neither is implied by the construction.

There is one free handle. The correlation data constrain only the off-diagonal structure of Σ^∗^. Its diagonal, the *S* marginal variances, is set by the noise amplitude and is not part of the data. The diagonal of Σ^∗^ therefore provides *S* free parameters, and it is exactly these *S* freedoms that we use to impose the *S* conditions 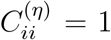. We solve these numerically and then test the resulting matrix for positive definiteness.

### 3. Sampling

We generate the synthetic data from a “true” interaction matrix whose off-diagonal entries are

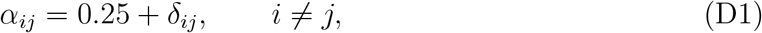

where *δ*_*ij*_ = *δ*_*ji*_ are Gaussian variables with standard deviation 0.09, and *α*_*ii*_ = 1. The environmental-noise correlation matrix is drawn independently, with

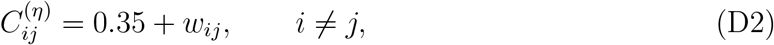

where *w*_*ij*_ = *w*_*ji*_ are Gaussian variables with standard deviation 0.10, and 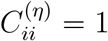. Draws are rejected unless both matrices are positive definite and the interaction matrix admits a positive coexistence fixed point. We take *S* = 6.

For each realization, the true interaction and noise matrices generate a target abundance correlation matrix. We then discard the true interaction matrix and ask whether the same abundance correlations can be explained by each of three deliberately incorrect candidates:

- an *independent* interaction matrix, drawn from the same ensemble and having the same mean overlap, but unrelated to the true matrix pair by pair.
- a *permuted* interaction matrix, obtained by shuffling the off-diagonal entries of the true matrix among species pairs. It therefore has exactly the same distribution of interaction strengths as the true matrix, but assigns those strengths to different pairs.
- a *sign-flipped* interaction matrix, *α*_*ij*_ ⟼ 2*µ*−*α*_*ij*_, which reverses the pairwise ordering of the interactions: the pairs with the strongest overlap in the true community are assigned the weakest overlap in the candidate, and conversely.

For each candidate, the *S* unknown marginal abundance variances are chosen so that the reconstructed environmental-noise matrix has unit diagonal, 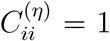 for *i* = 1, …, *S*. Once these variances are fixed, the Lyapunov construction reproduces the target abundance covariance identically. The only remaining nontrivial question is whether the reconstructed noise matrix is positive definite and can therefore represent a physically admissible environmental correlation structure.

**TABLE D1.** Very different interaction matrices generally require physically admissible environmental noise. Results for *S* = 6 over 200 independently generated communities. The second column is the mean Pearson correlation between the off-diagonal entries of the candidate and true interaction matrices. The third column counts the realizations for which the environmental-noise correlation matrix required by the candidate is positive definite. Its diagonal is fixed to unity by the choice of the marginal abundance variances. Every candidate reproduces the target abundance covariance exactly by construction. The table tests only whether the noise required to do so is a legitimate correlation matrix.

| Candidate interaction matrix | Mean correlation with the true interactions | Reconstructed noise positive definite |
| --- | --- | --- |
| Independent | +0.02 | 199/200 |
| Permuted | −0.03 | 200/200 |
| Sign-flipped | −1.00 | 200/200 |

### 4. Results

Table D1 summarizes 200 independent realizations. For every candidate that yields a positive-definite reconstructed noise matrix, the target abundance covariance is recovered exactly by construction. Numerically, the maximum reconstruction error was 2 × 10^*−*18^ for the independent and permuted candidates and 3 × 10^*−*18^ for the sign-flipped candidate, confirming the identity to machine precision.

The ambiguity is not confined to candidate matrices that resemble the true interactions. An independently drawn matrix requires a positive-definite noise correlation matrix in 199 of the 200 realizations, while both the permuted and sign-flipped candidates do so in every realization. Thus, even an interaction matrix that is unrelated, or exactly anticorrelated, with the true pairwise structure can reproduce the same abundance correlations using an ordinary, physically admissible environmental-noise correlation matrix. The non-identifiability is therefore not merely an algebraic consequence of allowing an arbitrary symmetric matrix on the right-hand side of the Lyapunov equation.

### 5. Behaviour with community size

Whether the degeneracy persists in larger communities must be asked at a fixed distance from the boundary of stability, not at fixed interaction heterogeneity. The spectral edge of the random part of ***α*** grows as 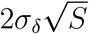, so holding *σ*_*δ*_ fixed while increasing *S* drives the community toward the instability threshold 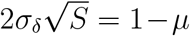. Near that threshold a single soft mode comes to dominate the covariance and stiffens the noise that can reproduce it. The controlling parameter is therefore not *S* but the reduced coupling 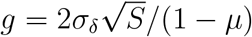 [38], the distance from criticality.

We repeated the admissibility test of the previous subsection along lines of constant *g*, scaling 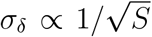 so that every community sits equally far from the stability boundary, and restricting the candidates to matrices that are themselves valid stable communities. At fixed *g* the admissible fraction does not fall with *S*: an independently drawn candidate remains admissible in essentially every realization, from a handful of species to several tens. The contraction that appears when *σ*_*δ*_ is instead held fixed is a near-critical effect, governed by *g* and setting in only as *g* → 1. It is not a property of diversity.

The degeneracy therefore does not close as the community grows. At any *S*, and at any fixed distance from criticality, the interaction matrix is not determined by the correlations alone. This is complementary to the localization result of Appendix E: the space of admissible interaction matrices stays open, and the correlations that any of them produces are governed by the centered mismatch 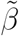, not by the niche overlap.

## Appendix E Abundance correlations for arbitrary *S*

The question addressed here is whether the environmental noise still governs the abundance correlations when the community is large, or whether its imprint is averaged away as the number of species grows. To settle it, this appendix derives Eq. (4), defines the quantity 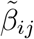 appearing in it, and gives the closed form valid at any community size. Throughout we work to leading order in the mismatch, using Eq. (4), in which the propagator is built from the homogeneous interaction matrix alone.

### 1. Solving the propagator equation

Let 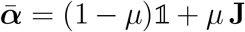, with **J** the matrix of ones, so that 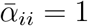 and 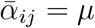 for *i* ≠ *j*. The fixed point is uniform,

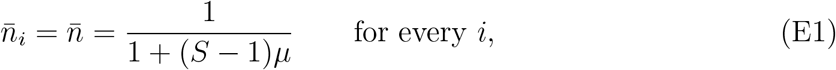

so 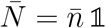 and 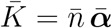 . Equation (22) then becomes

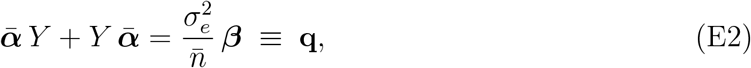

using the symmetry of 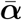. Substituting 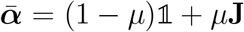 and writing *y*_*i*_ = ∑_*k*_ *Y*_*ik*_ for the row sums of *Y*, which is symmetric, we have (**J***Y* + *Y* **J**)_*ij*_ = *y*_*i*_ + *y*_*j*_, and Eq. (E2) reduces to a scalar relation for each pair,

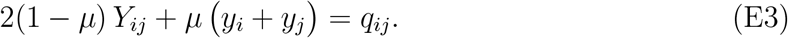

The *y*_*i*_ follow by summing. Writing *Q*_*i*_ = ∑_*j*_ *q*_*ij*_ and *Q* = ∑_*ij*_ *q*_*ij*_, and *Y* = ∑_*k*_ *y*_*k*_, one obtains

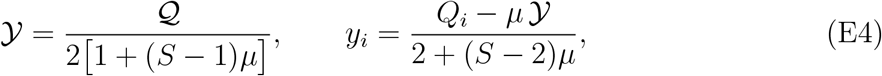

and then *Y*_*ij*_ from Eq. (E3). The symbols *y*_*i*_ and *Y* are internal to this step, whereas the quantities *c*_*i*_ introduced below in Eq. (E7) are row means of ***β*** and are a different object.

### 2. The correlations

Since 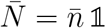, the ansatz (16) gives 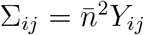 off the diagonal and 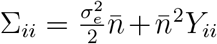 on it. To first order in ***β*** the second term of Σ_*ii*_ is negligible, and 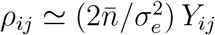. Substituting Eqs. (E3) and (E4), the factors of 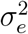 and 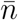 cancel, and we are left with a formula in ***β*** alone:

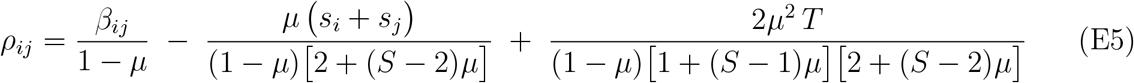

for *i* ≠ *j*, where

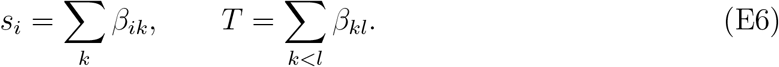

### 3. The large-*S* limit, and the definition of 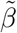

Equation (E5) has three terms, and they behave differently as the community grows. Introduce the row means and the grand mean of the mismatch,

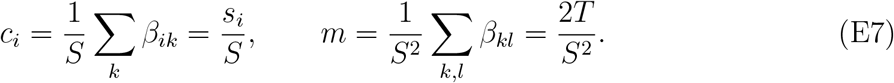

For *Sµ* ≫ 1 the denominators simplify, 2 + (*S* − 2)*µ* → *Sµ* and 1 + (*S* − 1)*µ* → *Sµ*, and the second and third terms of Eq. (E5) become (*c*_*i*_ + *c*_*j*_)*/*(1 − *µ*) and *m/*(1 − *µ*) respectively. Hence

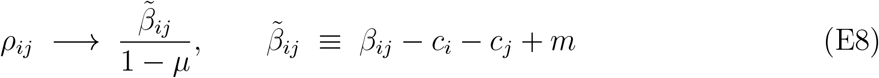

which is Eq. (4), and which defines 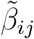: it is the mismatch of the pair (*i, j*) with the row, column and overall means removed. It is the residual of a two-way decomposition of ***β***, and it is what remains of the mismatch once everything that can be attributed to the individual species, or to the community as a whole, has been subtracted. The correlation therefore becomes *centered* at large *S*, not small: diversity removes the collective content of the mismatch, but 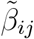 is an *O*(1) quantity, so *ρ*_*ij*_ does not vanish. The noise continues to set the correlations at every community size.

Numerically, for *S* = 60, *µ* = 0.2 and a homogeneous ***α*** (so that Eq. (E5) is exact rather than leading order, the heterogeneous case of Fig. 3 gives *r* = 0.98): the correlation between *ρ*_*ij*_ and 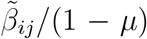 across the 1770 pairs is *r* = 0.9998, with slope 1.001. Against the *uncentred β*_*ij*_*/*(1 − *µ*) the agreement is visibly worse, *r* = 0.990 with slope 0.977, which is the signature of the collective terms that the centring removes.

The interpretation is the point of the exercise. Whatever part of the mismatch is shared by a species with all its competitors, or by the community as a whole, is suppressed: the first is absorbed into *c*_*i*_, the second into *m*, and both are divided out. What survives, and what a correlation measurement returns, is the part of the mismatch that is specific to *that pair*. Increasing diversity therefore does not average the ambiguity away. It localizes it: in a large community the correlation of a pair becomes a clean readout of that pair’s own mismatch, and of nothing else. Diversity sharpens the degeneracy rather than blurring it.

## Appendix F Adiabatic elimination of the resources

This appendix derives Eq. (4) of the main text. We take the environment to act on the resources alone, *σ*_*c*_ = 0 and *σ*_*r*_ *>* 0 in Eq. (24), and we eliminate the resources in the limit in which they are fast compared with the consumers.

Throughout the derivation we exploit the separation of timescales. On the fast resource timescale the consumer abundances are treated as frozen parameters: the resource fluctuations are solved for these fixed abundances, and only after averaging over the fast process is the resulting effective white noise reinserted into the slow consumer dynamics.

### 1. Fast resource fluctuations

Write 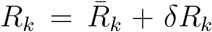, with 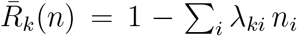 the quasi-steady level of Eq. (4). Here 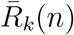 is the *instantaneous* quasi-steady resource level corresponding to the current consumer abundances, which are regarded as constant during the fast resource relaxation. On timescales short compared with the consumer dynamics the abundances may be held fixed, and linearizing Eq. (24b) about 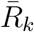 gives an Ornstein–Uhlenbeck process,

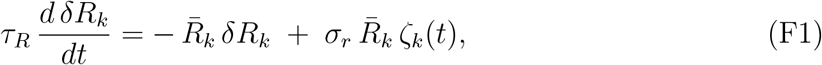

with relaxation rate and stationary variance

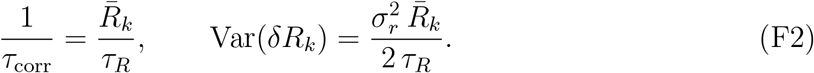

As *τ*_*R*_ is decreased the resource fluctuations become both faster and larger, and it is not obvious in advance which of the two effects wins.

### 2. Effective white-noise intensity

By Eq. (4), the growth rate of consumer *i* is displaced by ∑_*k*_ *γ*_*ik*_ *δR*_*k*_. The consumers are slow, so they do not resolve the shape of *δR*_*k*_(*t*), only its integrated effect: in the limit *τ*_*R*_ → 0 each *δR*_*k*_ acts on them as a white noise whose *intensity* is

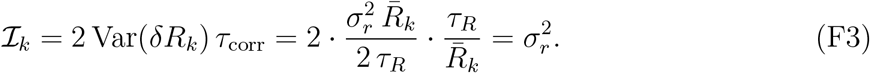

This is the central step, and it settles the question the two effects raised. The increasing variance and the decreasing correlation time cancel exactly, leaving a finite white-noise intensity independent of both *τ*_*R*_ and the equilibrium resource level. *No rescaling of the noise amplitude with the resource timescale is required*, and the weights are uniform across resources, 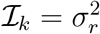 for every *k*.

### 3. The induced Lotka–Volterra description

Since the resources are uncorrelated, Eq. (4), the effective consumer-level noise 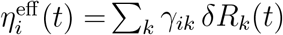 has covariance

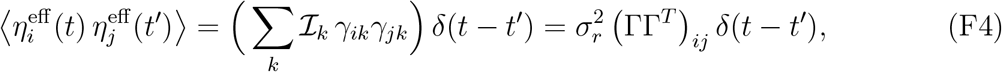

and the consumers obey a stochastic Lotka–Volterra system whose drift and diffusion matrices are

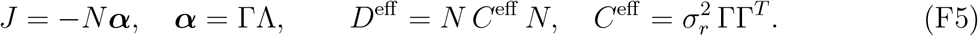

The coarse-graining therefore preserves the deterministic interaction matrix but replaces the environmental forcing by a different object. This separation is the mechanistic origin of the response–competition mismatch of the main text.

Two features distinguish *C*^eff^ from the Lotka–Volterra prescription of Sec. IV, and together they are the sources of the mismatch. First, competition is built from the yield *and* the depletion, whereas the inherited forcing is built from the yield *alone*: the reduction preserves only the product ΓΛ, so a given niche-overlap matrix is compatible with a continuum of factorizations, and hence with a continuum of mismatches ***β***^eff^ = *C*^eff^ − ***α***_*S*_ of either sign. Second, *C*^eff^ is not normalized to a unit diagonal: its diagonal entries 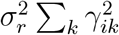 differ from species to species, so a yield generalist, spreading its yield over many resources, inherits less environmental variance than a yield specialist. The mismatch between *C*^eff^ and ***α*** therefore has two sources, the unequal diagonal amplitudes and the misalignment of yield and depletion, and both vanish only on the Λ ∝ Γ^*T*^ line, where the two matrices become proportional and the normalization *α*_*ii*_ = 1 renders the diagonal of ΓΓ^*T*^ uniform.

### 4. Numerical verification

The reduction is asymptotic, not exact, and must be checked. We integrated the full stochastic consumer–resource dynamics, Eq. (24) with *σ*_*c*_ = 0, directly in the Stratonovich convention, and compared the resulting pairwise abundance correlations with those of the reduced *S*-dimensional Lotka–Volterra system defined by Eq. (F5), holding *σ*_*r*_ *fixed* and applying no rescaling by *τ*_*R*_. The comparison is shown in Fig. F1. The two descriptions agree pair by pair, which confirms Eq. (F3): with *σ*_*r*_ held constant, and with no compensating rescaling, the reduced system reproduces the full dynamics in the fast-resource limit.

**FIG. F1.**
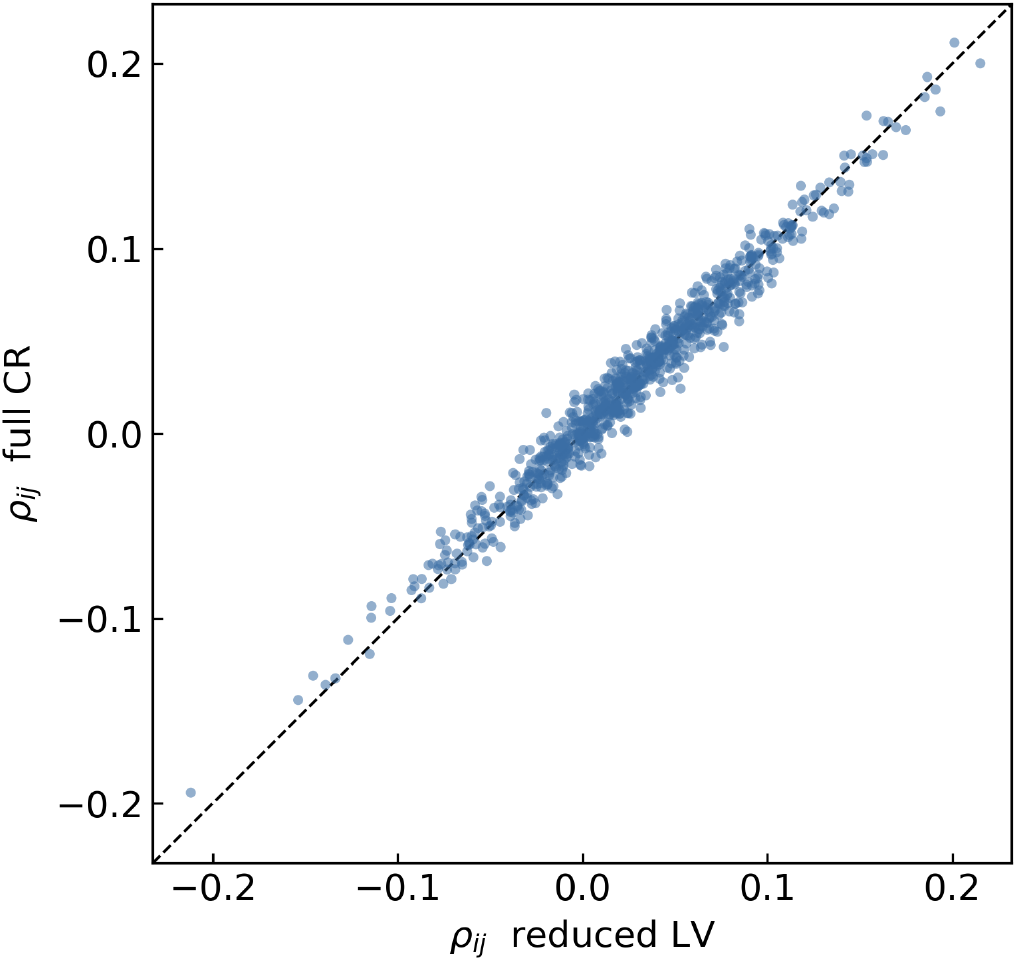
The reduction is exact in the fast-resource limit. Pairwise abundance correlations from direct Stratonovich simulation of the full stochastic consumer–resource system, Eq. (24) with *σ*_*c*_ = 0, against those of the reduced Lotka–Volterra system with ***α*** = ΓΛ and 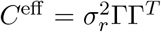. The dashed line is the identity, and *σ*_*r*_ is the same in both descriptions with no rescaling by *τ*_*R*_. Here *S* = 10 consumers and *Q* = 13 resources. Parameters *σ*_*r*_ = 0.02 and *τ*_*R*_ = 0.05. The plot shows the 45 pairs of each of 20 realizations. The Pearson correlation is 0.99 and the root mean square deviation from the identity is 0.01, the sampling error expected for a window of this length.

## Appendix G The yield–depletion mismatch

Section VI identifies the quantity that acts as a source of abundance correlations under resource-mediated stochasticity as the matrix

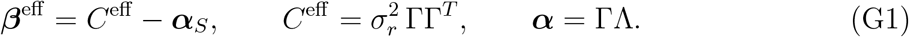

Still, Figure 5 of the main text is colored by the yield–depletion mismatch (YDM) of Ref. [16]. These two are quantities of different origin: ***β***^eff^ is the Lyapunov source that determines the covariance, and hence the correlations, whereas YDM was obtained by solving a simple two-species system exactly and guessing a similar-looking predictor of the correlations themselves. This appendix defines the YDM and gives its relation to ***β***^eff^, which is close but is not an identity.

### 1. Definition of Yield-depletion mismatch

Species *i* is characterized by two profiles across the resources: its yield *γ*_*i*_ = (*γ*_*i*1_, …, *γ*_*iQ*_), which says how much it gains from each resource, and its depletion *λ*_*i*_ = (*λ*_1*i*_, …, *λ*_*Qi*_), which says how much it removes. For a pair, the similarity of each profile is measured by its cosine, and the corresponding trait distance by

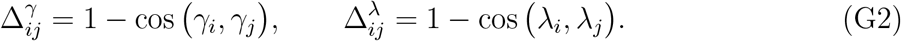

The yield–depletion mismatch is defined via the difference,

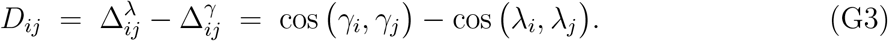

It is positive when two species draw their benefit from the same resources but deplete different ones, negative in the reverse case, and zero when the two profiles are equally similar. This is the convention of Ref. [16], in which differences in yield generate negative correlations and differences in depletion generate positive ones, so that *D* and *ρ*_*ij*_ carry the same sign.

### 2. Relation to *β*^eff^ : related, of different origin

***β***^eff^ and *D* are not the same object. ***β***^eff^ acts as the Lyapunov source, and as such it is an exact expression, but the correlations themselves reflect the value of ***β***^eff^ when filtered through the propagator of the Lyapunov relationships. *D* is instead a guess at those correlations, extracted from an exactly solvable two-species case [16].

To compare the yield–depletion mismatch with the exact dynamical mismatch ***β***^eff^, we revisit the exactly solvable two-consumer, two-resource example introduced in Appendix C of Ref. [16]. The yield and depletion matrices are chosen such that

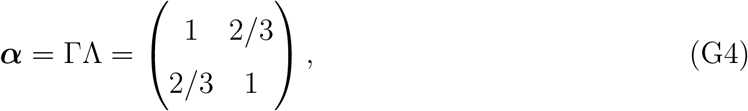

with

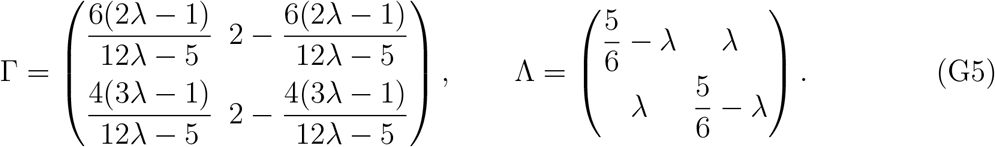

The exact abundance correlation is [16],

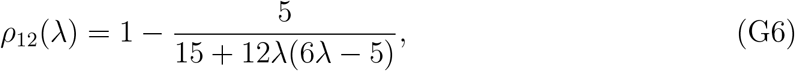

whereas the yield–depletion mismatch is

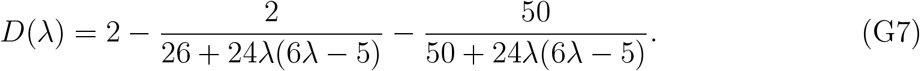

For resource-mediated forcing, let

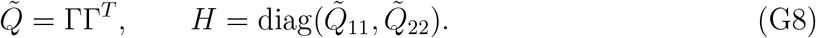

(We write 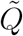 rather than *Q*, since *Q* denotes the number of resources throughout the paper.) The normalized weighted mismatch, in the convention ***β*** = *C*^(*η*)^ − ***α***_*S*_ of the main text (here *C*^eff^ against the matched noise 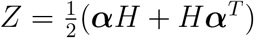 of Appendix C 3), is

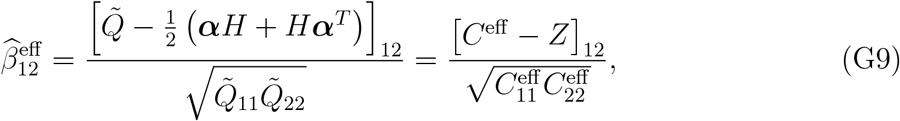

with 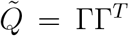 (so that 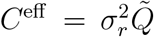) the noise actually inherited, and *Z* the noise that would annihilate the correlations. The common factor 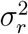 cancels between numerator and denominator, so 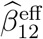 does not depend on the noise amplitude. For the matrices above this becomes

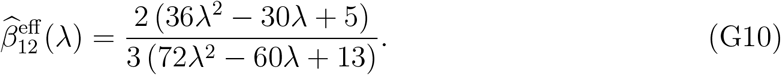

The three quantities are not identical, but they have the same sign and vanish at the same two values,

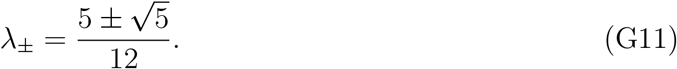

Thus, in this exactly solvable example, the geometrical proxy *D*(*λ*) and the weighted matrix mismatch 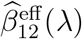 identify the same changes in the direction of the abundance correlation, although their magnitudes differ.

**FIG. G1.**
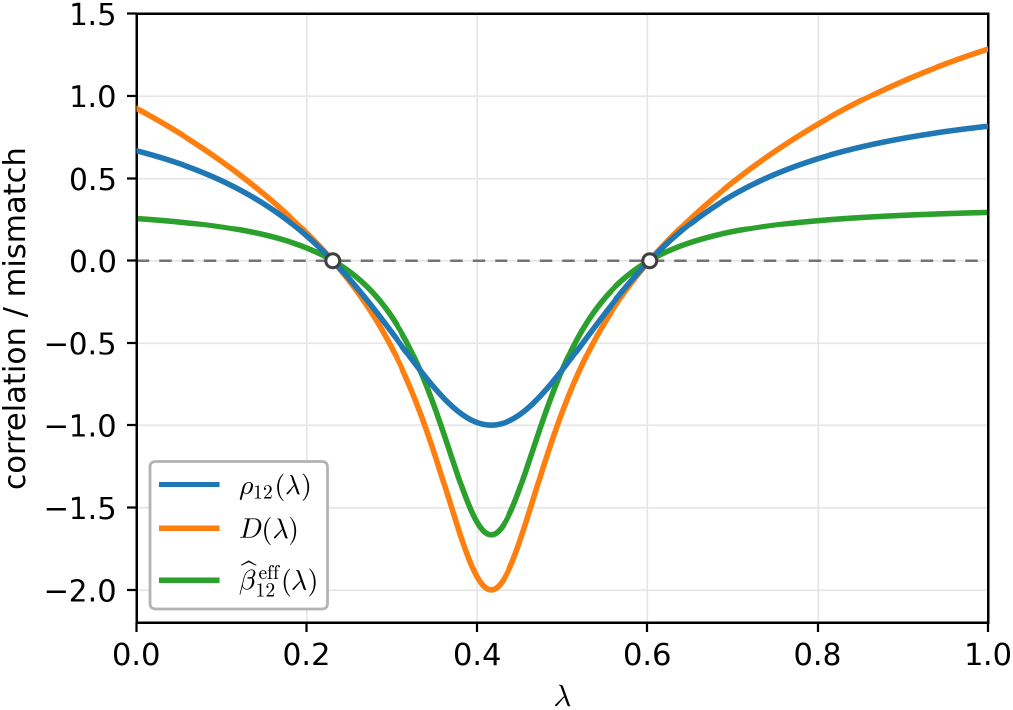
Exact correlation, yield–depletion mismatch, and weighted matrix mismatch in the solvable two-species example. The exact abundance correlation *ρ*_12_(*λ*) of Eq. (G6), the yield–depletion mismatch *D*(*λ*) of Eq. (G7), and the normalized weighted mismatch 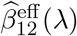 of Eq. (G10) are shown as functions of *λ*. The three quantities are not equal in magnitude, but have the same sign and cross zero at the same two values 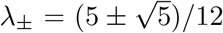, marked by the open circles. The exact Lyapunov mismatch and the geometrical YDM therefore encode the same qualitative transition in this analytically solvable case. At small and large *λ* the two consumers draw their benefit from overlapping resources while depleting different ones, so *D >* 0 and the abundances are positively correlated, and between *λ*_*−*_ and *λ*_+_ the relation is reversed.

